# The Individualized Neural Tuning Model: Precise and generalizable cartography of functional architecture in individual brains

**DOI:** 10.1101/2022.05.15.492022

**Authors:** Ma Feilong, Samuel A. Nastase, Guo Jiahui, Yaroslav O. Halchenko, M. Ida Gobbini, James V. Haxby

## Abstract

Quantifying how brain functional architecture differs from person to person is a key challenge in human neuroscience. Current individualized models of brain functional organization are based on brain regions and networks, limiting their use in studying fine-grained vertex-level differences. In this work, we present the Individualized Neural Tuning (INT) model, a fine-grained individualized model of brain functional organization. The INT model is designed to have vertex-level granularity, to capture both representational and topographic differences, and to model stimulus-general neural tuning. Through a series of analyses, we demonstrate that (a) our INT model provides a reliable individualized measure of fine-grained brain functional organization, (b) it accurately predicts individualized brain response patterns to new stimuli, and (c) it requires only 10–20 minutes of data for good performance. The high reliability, specificity, precision, and generalizability of our INT model affords new opportunities for building brain-based biomarkers based on naturalistic neuroimaging paradigms.

## 1. Introduction

A central goal of human neuroscience is to understand how brain functional organization differs across individuals, and how these differences relate to differences in intelligence, personality, motivation, mental health, and many other attributes. Understanding these differences is instrumental for providing individualized education and training, as well as effective diagnosis and intervention in the case of pathology, and ultimately improving educational, occupational, and health-related outcomes (Bijsterbosch et al., 2020; Dubois and Adolphs, 2016; Gabrieli et al., 2015; Gratton et al., 2020).

Models of the functional organization of the human brain can be summarized into two categories based on their spatial granularity. Typical functional magnetic resonance imaging (fMRI) data of the human brain comprises 20,000–100,000 cortical surface vertices (or voxels in volumetric data). Coarse-grained models group these vertices into spatial units—brain regions, networks, and systems—and reduce the brain into tens to hundreds of spatial units (Glasser et al., 2016; Gordon et al., 2016; Yeo et al., 2011). Vertices with similar, relatively homogeneous functions are studied as a group in coarse-grained models, which makes it easier to summarize their functions neuroscientifically and computationally (Bijsterbosch et al., 2020; Eickhoff et al., 2018b, 2018a). Recent advances of coarse-grained brain models have successfully extended group-level models to model individual brains (Gordon et al., 2017a; Harrison et al., 2015; Kong et al., 2019; Wang et al., 2015). In these models, the cortical topographies of the spatial units in an individual are allowed to differ from the group template, in order to account for inter-individual variations in brain functional organization (Gordon et al., 2017b; Gratton et al., 2018; Laumann et al., 2015). Individualized models help disentangle different sources of inter-individual variation (Bijsterbosch et al., 2019, 2018), and improve brain-behavior predictions (Kashyap et al., 2019; Kong et al., 2021).

Given this feature aggregation, coarse-grained models focus on spatial units that are centimeters in scale. Modern fMRI data acquisition, however, usually has a spatial resolution of 2–3 mm in each dimension, which is close to the spatial precision of blood-oxygen-level-dependent (BOLD) signal acquired at 3 Tesla (Engel et al., 1997; Parkes et al., 2005). This fine spatial resolution affords access to the rich information encoded in fine-grained vertex-by-vertex and voxel-by-voxel spatial patterns (Haxby et al., 2014, 2001; Huth et al., 2016; Kriegeskorte and Kievit, 2013). This information can be used to decode brain responses to different object categories (Haxby et al., 2001), and also different exemplars of the same category, such as different face identities or different views of the same face (Guntupalli et al., 2017; Visconti di Oleggio Castello et al., 2021, 2017). Individual differences in fine-grained responses and connectivity are much more reliable than their coarse-grained counterparts (Feilong et al., 2018). Fine-grained functional connectivity captures *what* information is exchanged between regions instead of *how much* information is exchanged, providing a twofold increase in accuracy in predicting intelligence (Feilong et al., 2021).

In this work, we present the individualized neural tuning (INT) model, a fine-grained individualized model of brain functional organization that has three key features. First, the INT model has vertex-level granularity, which provides access to the rich information encoded in fine-grained spatial patterns. Second, it models each individual’s unique representational geometry as well as the corresponding topographic organization in cortex, and thus affords study of both functional and topographic differences. Third, the INT model decomposes responses into stimulus information, as defined by neural responses that are shared across brains, and response tuning functions that model individual-specific fine-grained responses to any stimulus. Therefore, the INT model affords study of individual differences in neural response tuning that are independent of stimulus information (Figure 1).

**Figure 1.**
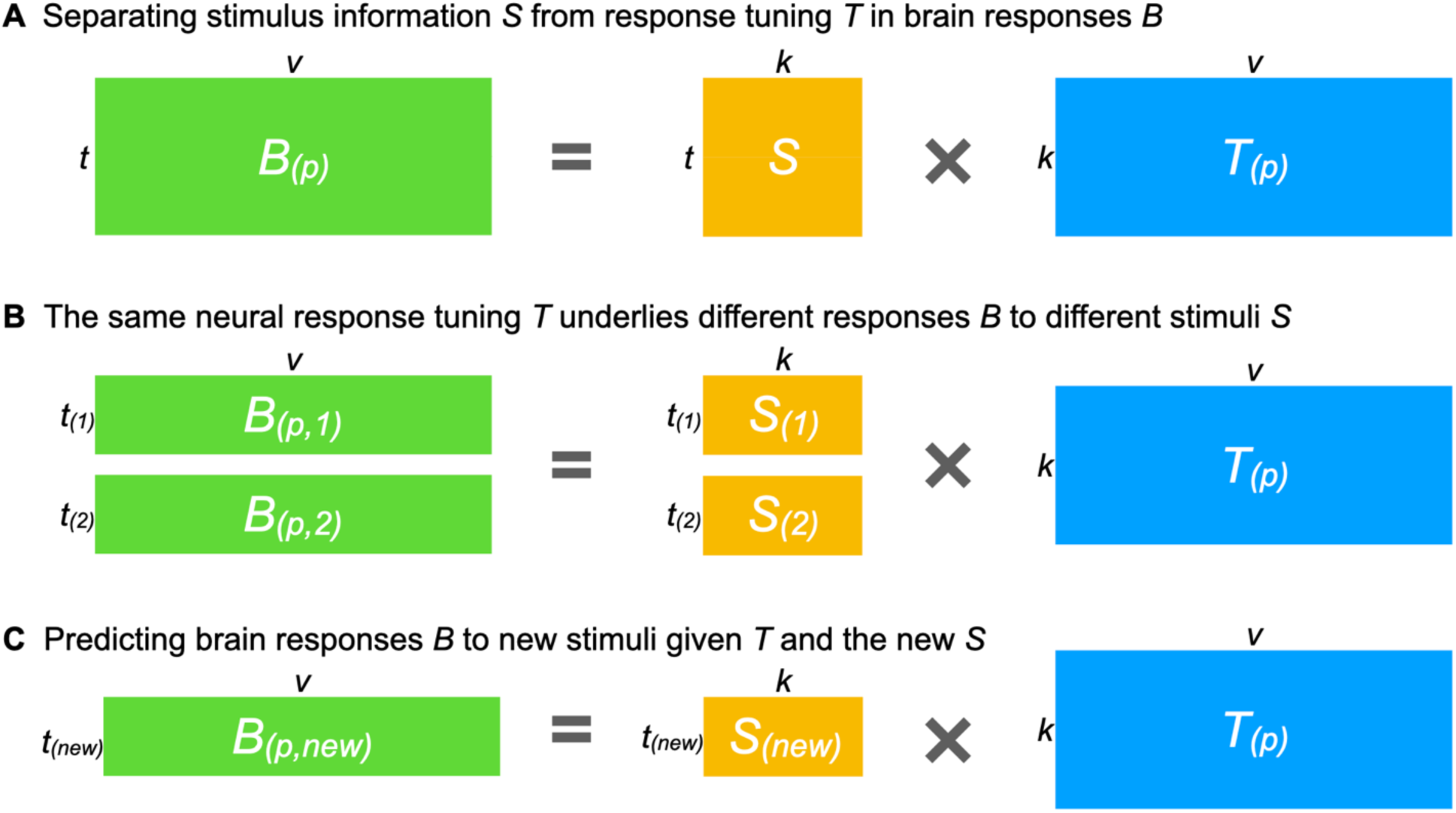
Estimating a shared stimulus matrix and individualized tuning matrices. **(A)** The individualized neural tuning (INT) model decomposes the brain response data matrix *B_(p)_* (shaped *t* × *v*, where *t* is the number of time points and *v* is the number of cortical vertices) of participant *p* into a shared stimulus matrix *S* (*t* × *k*, where *k* is the number of stimulus features) and an individualized tuning matrix *T_(p)_* (*k* × *v*, the number of stimulus features by the number of cortical vertices). Temporal information capturing how the stimulus changes over time is factored into *S*; each row of *S* is a time point in the stimulus and each column of *S* is a basis response profile shared across individuals and vertices. Each column of *T_(p)_* is a vector of *k* elements describing the response tuning function of a cortical vertex over basis response profiles. **(B)** If we divide the brain responses matrix *B_(p)_* into several parts (i.e., responses to different stimuli), each part can be modeled as part of the matrix *S* multiplied by the same *T_(p)_*. In other words, *T_(p)_* models neural response tuning in a way that generalizes across stimuli. Moreover, the same *T_(p)_* can be estimated from different parts of *B_(p)_* (e.g., two halves of a movie *B_(p,1)_* and *B_(p,2)_*) by using the corresponding parts of *S* (*S_(1)_* and *S_(2)_*). **(C)** After obtaining *T_(p)_*, it can be used to predict the participant’s responses to new stimuli *B_(p,new)_* using the corresponding *S_(new)_* matrix, which can be estimated from other participants’ data.

Using two rich fMRI datasets collected during movie watching, we demonstrate that our INT model of brain functional architecture has remarkable reliability and validity. Specifically, we show that: (a) two estimates of an individual’s model of brain function are highly similar based on independent data, but distinctive for different individuals; (b) the model can predict idiosyncratic patterns of brain responses to novel stimuli, including object categories and retinotopic localizers; (c) the model captures information encoded in fine-grained spatial patterns and can differentiate response patterns to different movie time points (TRs); and (d) the model works well with small amounts of movie data but continuously improves with more data. Together, these results demonstrate that our INT model predicts idiosyncratic fine-grained functional organization of the brain with high sensitivity and specificity.

## 2. Results

### 2.1. **Estimating the individualized neural tuning model**

Here we briefly describe the individualized neural tuning (INT) model in order to build a high-level intuition for how the model is constructed; see the “Methods” section for a more detailed mathematical treatment. Brain responses to external stimuli, such as movies, are broadly similar across individuals after anatomical alignment of cortical features and show much stronger similarity after the information contained in idiosyncratic fine-grained patterns is projected into a common model information space using hyperalignment (Guntupalli et al., 2018, 2016; Hasson et al., 2010, 2004; Haxby et al., 2020, 2011; Nastase et al., 2019). A substantial amount of an individual’s responses can be explained by these commonalities. Still, individuals differ from the common space and from each other, even though these differences are smaller in scale than the commonalities (Feilong et al., 2018). Therefore, it is critical to ensure that our model captures the idiosyncrasies of each individual’s brain functional organization, as well as the shared responses across individuals.

The goal of the INT model is to re-represent the brain data matrices *B_(p)_* acquired for each individual in a way that captures precise, individualized vertex-level functional architecture and supports out-of-sample prediction across both individuals and stimuli. First, we construct a common functional template *M* across all training participants to serve as a target for functional alignment based on all training participants’ data using a searchlight-based algorithm. Next, we estimate a linear transformation *W_(p)_* for each participant, using ensemble ridge regression, that maps between their idiosyncratic functional architecture and the functional template *M*. Unlike previous implementations of hyperalignment that employed Procrustes-based rotations to resolve topographic idiosyncrasies while preserving representational geometry, here we estimate a linear transformation that captures individual differences in both representational geometry and cortical topography. Finally, we convert the model-estimated brain data, *MW_(p)_*, into a more compact shared stimulus matrix *S,* with orthogonal feature dimensions, and an individualized tuning matrix *T_(p)_*. This decomposition factors the stimulus-specific temporal structure of the movie into *S*, represented as a collection of basis functional profiles shared across vertices and individuals. The individual-specific tuning matrices *T_(p)_* can be estimated with independent data using different stimuli. The *T_(p)_* matrices capture individual differences in functional tuning—modeling idiosyncrasies in both representational geometry and cortical topography.

### 2.2. Modeling individualized brain functional organization

To assess how well our model captures individual-specific brain functional organization, we evaluated the within-subject similarities and between-subject similarities of the modeled tuning matrices (*T*). For each of the *n* participants, we divided the movie data into two parts, and computed a tuning matrix independently for each movie part. Therefore, we obtained estimates of *n* tuning matrices based on the first part of the movie, and an independent set of *n* estimated tuning matrices based on the second part. Then we computed an *n* × *n* matrix of cross-movie-part similarities, where each row corresponds to a tuning matrix based on the first part, and each column corresponds to a tuning matrix based on the second part. Each entry in the matrix quantifies the cross-movie-part similarity of tuning matrices within-subject (diagonal entries) and between-subject (off-diagonal entries) (Figure 2A). For both datasets, the similarity matrix had a clear diagonal, indicating that the within-subject similarities were much higher than between-subject similarities. When all the tuning matrices were projected to a 2-D plane using multi-dimensional scaling (MDS), matrices from the same participant were close together, whereas matrices from different participants were clearly separated (Figure 2B).

**Figure 2.**
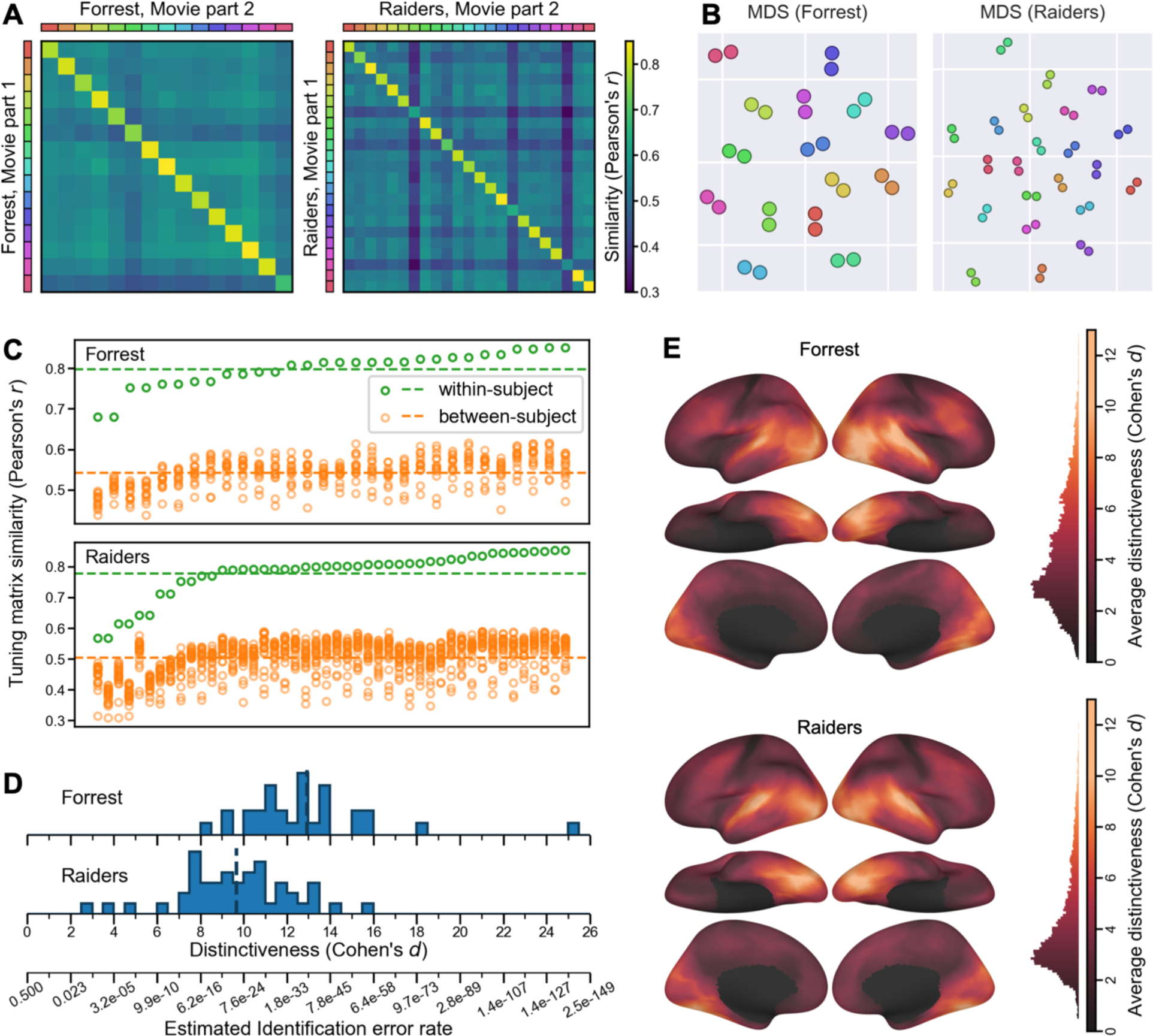
Modeling individual-specific brain functional organization. **(A)** For each movie part, we obtained *n* tuning matrices, one for each participant, which describes the participant’s response tuning functions. The cross-movie-part similarities form an *n* × *n* matrix, where rows are tuning matrices based on the first movie part, and columns the second movie part; the colored legends at left and top index individual participants. The obvious diagonal indicates that within-subject similarities were much higher than between-subject similarities. **(B)** Multi-dimensional scaling (MDS) projection of the 2*n* matrices onto a 2-D plane. Two dots of the same color denote two estimates of the tuning matrix for the same participant, as in **(A)**. Dots from the same participant clustered together. **(C)** The distribution of within- and between-subject tuning matrix similarities, sorted by within-subject similarity. For each tuning matrix, the within-subject similarity always exceeded between-subject similarity. **(D)** We computed a distinctiveness index for each tuning matrix based on the difference between within- and between-subject similarities. The distinctiveness index is based on Cohen’s *d* and, therefore, measures effect size. Based on the distinctiveness index, we estimate the error rate for individual identification (bottom). **(E)** Local functional distinctiveness based on a searchlight analysis (20 mm radius), averaged across all participants for each dataset. Extensive occipital, temporal, and lateral prefrontal cortices showed high distinctiveness.

For every tuning matrix, within-subject similarities (*Forrest*: *r* = 0.798 ± 0.044 [mean ± SD]; *Raiders*: *r* = 0.778 ± 0.076) were higher than between-subject similarities (*Forrest*: *r* = 0.542 ± 0.037; *Raiders*: *r* = 0.503 ± 0.057) (Figure 2C). Simple nearest-neighbor identification of participants based on their tuning matrices performs at 100% accuracy. To better assess the distinctiveness of each tuning matrix, we computed a distinctiveness index based on Cohen’s *d* (Figure 2D). This distinctiveness index measures the difference between the within-subject similarity and between-subject similarities of a tuning matrix using the standard deviation of the distribution as a unit. For example, Cohen’s *d* = 5 means that the within-subject similarity is 5 standard deviations greater than the average between-subject similarity. On average across participants, the distinctiveness index was 12.92 for the *Forrest* dataset, and 9.67 for the *Raiders* dataset, indicating the individual-specific tuning matrices were highly distinctive. The distinctiveness index was computed based on Fisher-transformed correlation similarities, which approximately follow a normal distribution. Therefore, the identification error rate can be estimated based on the distinctiveness index using the cumulative distribution function of the distribution, which was 1.73×10^-38^ for *d* = 12.92, and 2.1×10^-22^ for *d* = 9.67. These error rates are orders of magnitude lower than those estimated from individuation based on coarse-grained patterns of functional connectivity (more than 1%; (Finn et al., 2015)) and those of forensic DNA analysis (approximately 0.4%; (Kloosterman et al., 2014)).

The results so far are based on the entire tuning matrix, which comprises response tuning functions of all cortical vertices. Which part of the brain has the most distinctive responses across individuals? To answer the question, we performed a searchlight analysis with a 20 mm radius and computed the average distinctiveness index across participants for each searchlight (Figure 2E). Extensive occipital, temporal, and lateral prefrontal cortices showed high distinctiveness, with estimates of Cohen’s *d* exceeding 10 in lateral and ventral occipital and temporal cortices. Even in brain regions that do not respond strongly to external stimuli, such as medial prefrontal cortex, our model can still capture idiosyncratic response tuning functions. To summarize, our model of brain functional organization is highly specific to each individual. For both datasets, within-subject similarities of modeled tuning matrices were several standard deviations higher than between-subject similarities. Our model also captures idiosyncrasies in local response tuning functions throughout cortex, excluding somatosensory and motor regions. Individual differences were most prominent in occipital and temporal regions, and reliable individual differences were also found in parietal and prefrontal regions.

### 2.3. Predicting category-selectivity and retinotopic maps

To assess whether the modeled tuning matrix accurately reflects a participant’s brain functional organization, we examined to what extent it can predict brain responses to new stimuli. Specifically, we examined whether our model trained with movie data could accurately predict category-selectivity maps and retinotopic maps in a leave-one-subject-out cross-validation analysis.

#### 2.3.1. Predicting category-selectivity maps

Both the *Forrest* dataset and the *Raiders* dataset had four object category localizer runs, which were based on static images for *Forrest*, and dynamic videos for *Raiders*. Taking the “faces” category as an example, we computed a face-selectivity map for each participant and each run, which was the contrast between faces and all other categories. Due to measurement noise, the four maps generated for each individual participant (one for each run) differ from one another (Figure 3B and 3C bottom rows). We averaged the four maps for each participant to reduce noise and used the average map as the localizer-based map for that participant. Based on the similarity between these four maps, we computed the Cronbach’s alpha coefficient for each participant, which estimates the reliability of the average map. That is, if we were to scan the participant for another four localizer runs and correlate the new average map with the current average map, the expected correlation would be Cronbach’s alpha.

**Figure 3.**
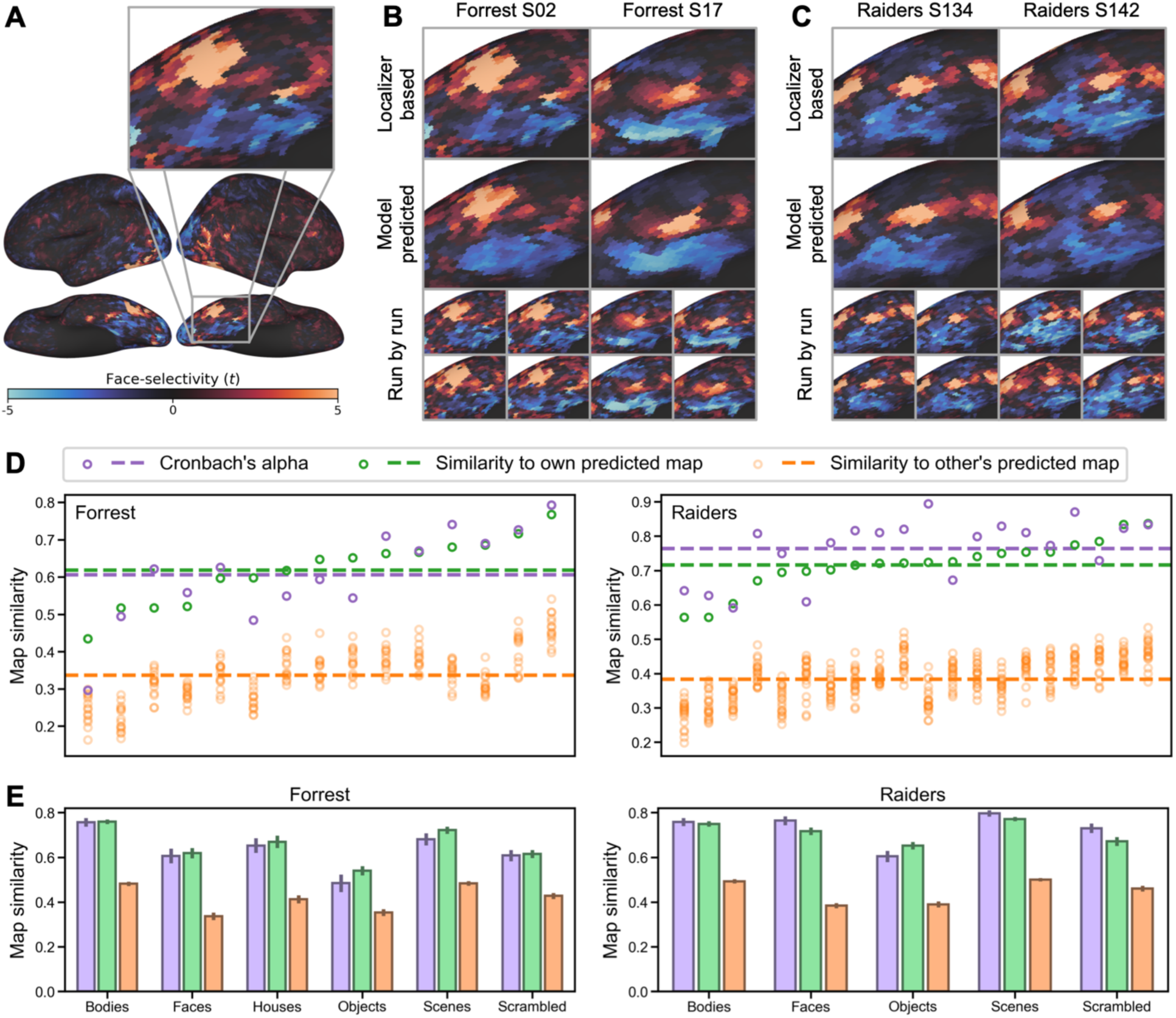
Predicting category-selectivity maps of individual participants. **(A)** Face-selectivity map of an example participant and a zoomed-in view focusing on right ventral temporal cortex. **(B)** The localizer-based (top) and model-predicted (middle) face-selectivity maps for two example participants from the *Forrest* dataset. Each localizer-based map was the average of four maps, one from each localizer run. Individual maps for each localizer run are shown at bottom. **(C)** Face-selectivity maps of two example participants from the *Raiders* dataset. **(D)** Similarity of each participant’s localizer-based face-selectivity map to the participant’s own predicted map (green) and to other participants’ predicted maps (orange). Cronbach’s alpha (purple) for each participant was calculated based on the similarity of the four localizer runs and is shown as a reference. **(E)** Cronbach’s alpha (purple), within-subject correlation (green), and between-subject correlation (orange) for all category-selectivity maps. Error bars are standard errors of the mean. For both datasets, the within-subject correlations were similar to, and sometimes higher than Cronbach’s alpha. Between-subject correlations were much lower, suggesting our prediction models were able to capture each participant’s idiosyncratic category-selectivity topographies.

For each cross-validation fold, we divide the data into *n* – 1 training participants and a test participant. To estimate the stimulus descriptors for the target object category (e.g., *S*_(faces)_), we trained a regression model to predict the localizer-based maps for the training participants (dependent variables) from their tuning matrices (*T*) (independent variables). The resultant *S*_(faces)_ vector contains the coefficients derived from the regression model. *T* was estimated from the independent movie data for each participant and applied to this analysis. Then we computed the product of the *S*_(faces)_ vector of coefficients and the test participant’s tuning matrix (*T*) to estimate the test participant’s face-selectivity map. We evaluated the quality of this predicted localizer map by computing the correlation between the model-based map and the test participant’s actual localizer map based on their own localizer data.

For both datasets, the localizer-based and model-predicted face-selectivity maps were highly correlated (*Forrest*: *r* = 0.618 ± 0.089 [mean ± SD], *Raiders*: *r* = 0.716 ± 0.074), and the correlations were higher than our previous state-of-the-art hyperalignment model with the same dataset (Jiahui et al., 2020). Across all participants, the average Cronbach’s alpha was 0.606 ± 0.126 for *Forrest*, and 0.764 ± 0.089 for *Raiders*. For approximately a third of the participants (*Forrest*: 6 out of 15, 40%; *Raiders*: 6 out of 20, 30%), the correlation exceeded the Cronbach’s alpha of localizer-based maps. In other words, for these participants, the predicted maps based on our model were more accurate than the maps based on a typical localizer scanning session comprising four runs.

Besides the high accuracy, the model-predicted maps were also highly specific for each individual (See Figure 3B and 3C for examples). The correlation between one participant’s localizer-based map and another participant’s model-predicted map (orange circles in Figure 3D; *Forrest*: 0.337 ± 0.071; *Raiders*: 0.384 ± 0.062) was always lower than the correlation with own model-predicted map (green circles in Figure 3D). This indicates that our model accurately predicts the idiosyncratic topographies of each participant’s category-selectivity map.

We replicated our analysis for all other categories and found similar results (Figure 3E; Table 1). For all object categories and both datasets, the within-subject similarity (correlation between own localizer-based map and own model-predicted map) was numerically similar to Cronbach’s alpha and much larger than between-subject similarities (correlation between each participant’s localizer-based map and others’ model-predicted maps).

**Table 1.** Summary of model performance in predicting object category selectivity maps. All contrasts were based on the target category versus all others. The format for Cronbach’s alpha and similarities is mean ± standard deviation.

| The *Forrest* dataset |  |  |  |  |  |
| --- | --- | --- | --- | --- | --- |
| Category | Cronbach's alpha | Within-subject similarity | Between-subject similarity | (within > between) % | (within > alpha) % |
| Bodies | $0.756 \pm 0.073$ | $0.759 \pm 0.041$ | $0.482 \pm 0.037$ | 100% | 40.0% |

**Table 1. Summary of model performance in predicting object category selectivity maps.** All contrasts were based on the target category versus all others. The format for Cronbach's alpha and similarities is mean ± standard deviation.
| Faces | 0.606 ± 0.126 | 0.618 ± 0.089 | 0.337 ± 0.064 | 100% | 40.0% |
| --- | --- | --- | --- | --- | --- |
| Houses | 0.653 ± 0.128 | 0.669 ± 0.106 | 0.412 ± 0.070 | 100% | 46.7% |
| Objects | 0.485 ± 0.153 | 0.540 ± 0.079 | 0.353 ± 0.058 | 100% | 60.0% |
| Scenes | 0.681 ± 0.107 | 0.721 ± 0.063 | 0.483 ± 0.040 | 100% | 53.3% |
| Scrambled | 0.608 ± 0.096 | 0.615 ± 0.070 | 0.427 ± 0.051 | 100% | 60.0% |
| The *Raiders* dataset |  |  |  |  |  |
| Category | Cronbach's alpha | Within-subject similarity | Between-subject similarity | (within > between) % | (within > alpha) % |
| Bodies | 0.758 ± 0.083 | 0.749 ± 0.056 | 0.493 ± 0.042 | 100% | 45.0% |
| Faces | 0.764 ± 0.089 | 0.716 ± 0.074 | 0.384 ± 0.051 | 100% | 30.0% |
| Objects | 0.604 ± 0.113 | 0.652 ± 0.077 | 0.390 ± 0.061 | 100% | 65.0% |
| Scenes | 0.796 ± 0.061 | 0.771 ± 0.043 | 0.500 ± 0.034 | 100% | 30.0% |
| Scrambled | 0.730 ± 0.096 | 0.671 ± 0.089 | 0.461 ± 0.057 | 100% | 40.0% |

#### 2.3.2. Predicting retinotopic maps

We examined whether our model can accurately predict eccentricity and polar angle maps based on the retinotopic data of the *Forrest* dataset. Similar to category-selectivity maps, we trained our model using the movie data and used it to predict retinotopic maps based on leave-one-subject-out cross-validation. Note that each retinotopic map, eccentricity and polar angle, has two components: an amplitude map, which measures to what extent a cortical vertex responds to retinotopic stimuli, and a phase map, where the phase is associated with eccentricity or polar angle. For the eccentricity map, the phase is 0° for the center of the visual field, and 360° for the most peripheral part. For the polar angle map, the phase is 0° and 180° for the upper and lower vertical meridians, and 90° and 270° for the right and left horizontal meridians.

The model-predicted maps for each participant resemble the corresponding localizer-based maps, and they capture the idiosyncratic features of each map well (Figure 4A and 4B). To quantify these similarities, we assessed the similarity of amplitude maps and phase maps separately. Each retinotopic map (e.g., an eccentricity map) was based on a standard univariate analysis of two runs where the stimuli were displayed in reversed order (e.g., expanding rings and contracting rings), and an amplitude map and a phase map were obtained from each run. For each participant, we compared the similarity of these two amplitude maps and estimated Cronbach’s alpha. The mean (± standard deviation) for Cronbach’s alpha was 0.701 ± 0.047 for the eccentricity map, and 0.663 ± 0.069 for the polar angle map. We also compared the similarity between the localizer-based amplitude map (average of the two runs) and the model-predicted map. On average across all participants, the similarity was 0.774 ± 0.027 for the eccentricity map, and 0.746 ± 0.049 for the polar angle map. Note that for every participant the similarity was higher than Cronbach’s alpha, which means the model-predicted amplitude map is more accurate than the localizer-based map. The similarity between a participant’s localizer-based map with the participant’s own model-predicted map is higher than with others’ model-predicted maps (eccentricity: 0.682 ± 0.029; polar angle: 0.635 ± 0.054), indicating that the model-predicted amplitude map is individual-specific.

**Figure 4.**
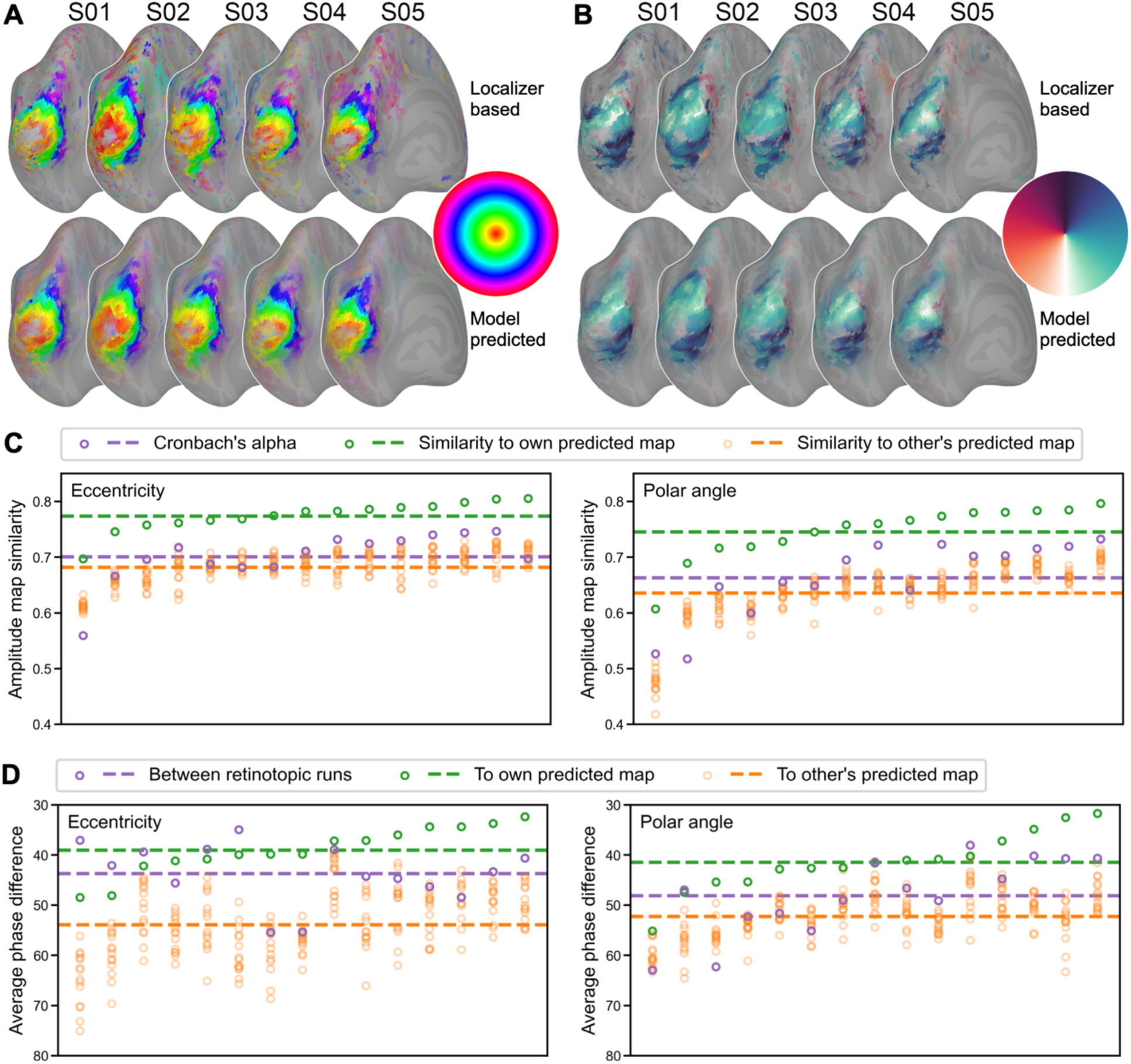
Predicting retinotopic maps of individual participants. **(A)** The localizer-based (upper) and model-predicted (lower) left hemisphere eccentricity and **(B)** polar angle maps for five example participants. **(C)** Similarity of each participant’s localizer-based amplitude map (i.e., to what extent a vertex responds to retinotopic stimuli) to the participant’s own predicted map (green), other participants’ predicted maps (orange), and its Cronbach’s alpha (purple). **(D)** The average phase difference in early visual areas between the participant’s two retinotopic runs (e.g., expanding and contracting rings; purple), between the participant’s localizer-based map and own model-predicted map (green), and between the participant’s localizer-based map and other participants’ predicted maps (orange). In both **(C)** and **(D)** participants are sorted along the x-axis according to within-subject similarity (green). Note that we inverted the y-axis in **(D)** because smaller differences indicate higher similarity.

To assess the quality of the phase maps, we computed the absolute value of the phase difference in early visual areas (V1, V2, V3, and V4; (Glasser et al., 2016)) between two retinotopic runs, between the localizer-based map and the participant’s own model-predicted map, and between one participant’s localizer-based map and others’ model-predicted maps. Note that the phase is circular, and thus the difference between 360° and 1° is the same as 1° and 2°. On average across participants, the average phase difference between a participant’s localizer-based and model-predicted maps was 39.1° ± 4.8° for eccentricity maps, and 41.5° ± 6.0° for polar angle maps. This difference was smaller than the difference between two localizer runs (eccentricity: 43.7° ± 6.0°; polar angle: 48.2° ± 7.7°) and the difference with others’ model-predicted maps (eccentricity: 53.9° ± 6.9°; polar angle: 52.3° ± 4.7°). The average phase difference for random data would be 90°.

For both category-selectivity maps and retinotopic maps, our model can accurately predict individualized maps with high fidelity and high specificity. The quality of the model-predicted maps was similar to or higher than that of maps derived from actual localizer data. These results demonstrate that the modeled response tuning functions are not only individualized and reliable across independent data, but also can accurately predict responses to new stimuli.

### 2.4. Predicting brain responses to the movie

The previous analyses show that our model accurately predicts brain responses for category-selectivity and retinotopic maps. These maps reflect coarse-grained functional topographies of the brain: they are relatively spatially smooth, and neighboring vertices on the cortex (especially vertices in the same brain region) have similar category-selectivity or adjacent receptive fields. In the analysis below, we examine whether our model can accurately predict fine-grained functional topographies; that is, the vertex-by-vertex spatial patterns which vary substantially even within a brain region. Rich visual, auditory, and social information is encoded in fine-grained spatial patterns of response (Haxby et al., 2014). Specifically, we trained our model using half of the movie data and predicted the other half.

We used a leave-one-subject-out cross-validation to evaluate the performance of our INT model. We derived the tuning matrix *T* of the test participant based on the first half of the participant’s movie data, and combined it with *S_(2)_* (the part of *S* for the second part of the movie, derived from the training participants’ data) to predict the test participant’s responses to the second part of the movie. The response pattern at each time point (i.e., TR) of the movie comprises 18,742 values, one for each cortical vertex. Similar to our previous work (Guntupalli et al., 2016), we trained a principal component analysis (PCA) based on the first half of the movie to reduce dimensionality from 18,742 vertices to a few hundred principal components (PCs) and projected responses to the other half of the movie onto these PCs. Analysis of whole-brain spatial patterns of response was based on these normalized PCs.

The model-predicted response patterns for the movie were highly specific to both the time point and the participant. Note that these model-predicted patterns are based on other participants’ neural responses projected into the native, fine-grained cortical topography of the left-out test participant’s brain. The predicted pattern for a certain time point for a left-out test participant’s brain was much more similar to the measured response pattern to the same time point in that participant’s brain (Figure 5A diagonal) than responses to other time points (Figure 5A off-diagonal). The average correlation similarity between predicted and measured response patterns for the same time point was 0.356 for the *Forrest* dataset, and 0.408 for the *Raiders* dataset, whereas the average similarity between predicted and measured patterns from different time points was close to 0 for both datasets. For the same time point, the measured response patterns were more similar to predicted patterns in a participant’s native space than to predicted patterns in other participants’ native spaces (Figure 5B diagonal). The average similarity of the same time point for different participants was 0.211 for the *Forrest* dataset, and 0.209 for the *Raiders* dataset (Figure 5C).

**Figure 5.**
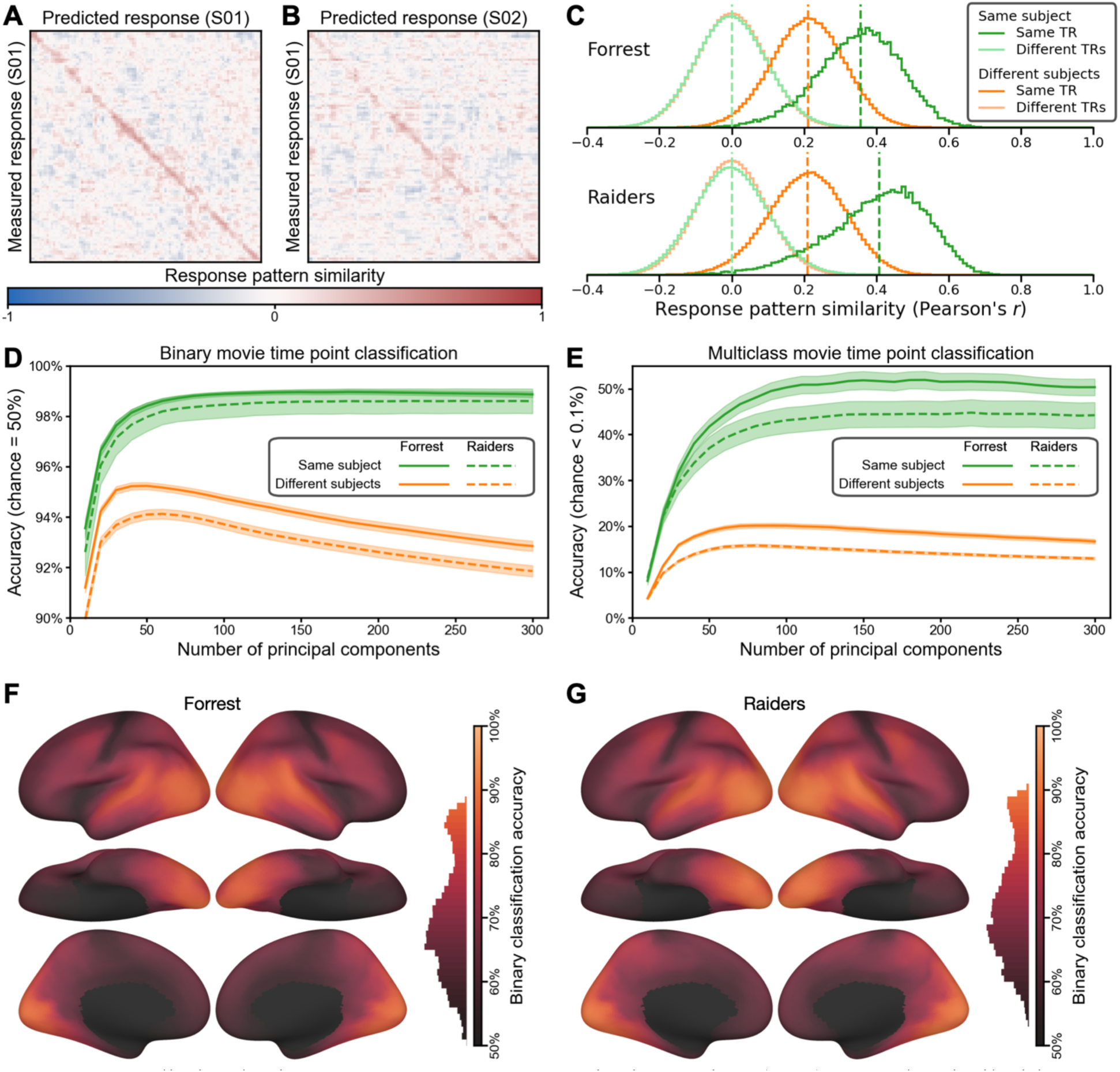
Predicting brain response patterns to movie time points (TRs). **(A)** The similarities between measured and predicted brain response patterns for the first 100 time points of an example *Forrest* participant (the full matrices for *Forrest* and *Raiders* contain 1818 and 1680 time points, respectively). The red diagonal indicates that the model-predicted response pattern at each time point was highly similar to the actual response pattern for the corresponding time point. The response patterns were based on 150 principal components (PCs) reduced from all cortical vertices. **(B)** The similarities between measured response patterns of one participant and predicted patterns of another. The less obvious diagonal suggests that our model predicted both the shared functional topographies (which generalize across participants) and each participant’s idiosyncratic functional topographies (which does not generalize across participants). **(C)** The distribution of response pattern similarities across participants and time points. When the measured and the predicted patterns were for the same time point of the movie, the average within- and between-subject similarities were 0.356 and 0.211, respectively, for the *Forrest* dataset, and 0.408 and 0.209, respectively, for the *Raiders* dataset. Cross-time-point similarities were centered around 0. This indicates that the predicted movie response patterns were highly specific to both the participant and the time point. **(D)** Binary (2-alternative forced choice) movie time point classification based on a nearest-neighbor classifier and pattern similarities. The within-subject accuracy peaked at 99.0% for *Forrest* (180 PCs) and 98.6% for *Raiders* (250 PCs), and it was fairly robust across the number of PCs. The peak between-subject accuracy was 95.2% (50 PCs) and 94.1% (60 PCs), respectively. **(E)** Multiclass movie time point classification. The number of choices was 1818 for *Forrest* and 1680 for *Raiders*, and chance accuracy was less than 0.1% for both datasets. The peak within-subject accuracy was 51.9% for *Forrest* (190 PCs) and 44.8% for *Raiders* (220 PCs), and the peak between-subject accuracy was 20.1% for *Forrest* (90 PCs) and 15.8% for *Raiders* (80 PCs). **(F and G)** Searchlight binary classification. The accuracy was high for much of the cortex for both datasets, with the highest accuracies in temporal and occipital regions.

Considering the similarity between measured and predicted response patterns, we assessed whether we could classify which time point of the movie the participant was viewing based on these patterns. We performed the classification analysis using a one-nearest-neighbor classifier in two different ways. First, we used binary classification (2-alternative forced choice); that is, we compared the measured response pattern for one time point with the predicted patterns for the same single time point paired with each other time point to determine which pair is more similar, and then averaged across all pairs, resulting in a chance accuracy of 50%. Second, we used multiclass classification; that is, whether the similarity with the same time point is higher than with all other time points. The number of time points was 1818 for *Forrest* and 1680 for *Raiders*, resulting in a multiclass chance accuracy less than 0.1% for both datasets. We varied the number of PCs used in the analysis from 10 to 300 with an increment of 10 and repeated the analysis at each number of PCs. For binary classification, the accuracy peaked at 99.0% for *Forrest* (180 PCs) and 98.6% for *Raiders* (250 PCs) (Figure 5D). For multiclass classification, the peak accuracy was 51.9% for *Forrest* (190 PCs) and 44.8% for *Raiders* (220 PCs) (Figure 5E). Note that these classification results are robust against the number of PCs used, and the accuracy was stable with 100–300 PCs for both approaches and both datasets.

The response patterns of different participants’ share some similarities (Figure 5C, dark orange), and we were able to classify which time point one participant was viewing based on the predicted patterns in another participants’ native space to some extent. For the binary classification analysis, the peak accuracy was 95.2% for *Forrest* (50 PCs) and 94.1% for *Raiders* (60 PCs) (Figure 5D, orange lines). For the multiclass classification analysis, the peak accuracy was 20.1% for *Forrest* (90 PCs) and 15.8% for *Raiders* (80 PCs) (Figure 5E, orange lines). Note that the classification accuracy for mismatching participants drops dramatically after peaking at 50–90 PCs, whereas the classification accuracy for the matching participant monotonically improved until the number of PCs is roughly 200. This suggests that a considerable amount of the information in our model-predicted response patterns are specific to the test participant. To localize cortical areas where the fine-grained patterns are most accurately predicted, we performed a searchlight analysis (20 mm radius) with the binary classification approach. Due to the limited number of vertices in each searchlight, we performed the classification analysis without dimensionality reduction. We found that the accuracy was highest for visual, auditory, and corresponding association cortices (Figure 5F & G) with significant classification across almost all of cortex.

### 2.5. Model performance with less data

The datasets used so far in this work comprise relatively long-duration movie-watching fMRI acquisitions (*Forrest*: 120 minutes; *Raiders*: 56 minutes), which may not be feasible for every fMRI experiment due to limited scanning resources. How well does our INT model work with smaller amounts of movie data? To address the question, we systematically manipulated the amount of movie data for the test participant and assessed our model performance for key benchmarking indices. For the *Forrest* dataset, the durations were 5, 10, 15, 20, 30, 40, 50, 60, and 120 minutes; for the *Raiders* dataset, the durations were 5, 10, 15, 20, 28, and 56 minutes. Depending on the analysis, up to half of the movie data (60 and 28 minutes, respectively) or the entire movie dataset was used.

With more movie data used for estimating a tuning matrix, the distinctiveness of that modeled tuning matrix increased monotonically (Figure 6A). With 10 minutes or more movie data, the average Cohen’s *d* was more than 6, which means within-subject similarity of tuning matrices exceeded between-subject similarities by more than six standard deviations on average. Given that Fisher-transformed correlation similarities are approximately normally distributed, the chance of a between-subject similarity exceeding the within-subject similarity was less than 10^-9^. In other words, if we were to identify an average individual using the tuning matrix based on 10 minutes of movie data, the error rate would be less than 10^-9^.

**Figure 6.**
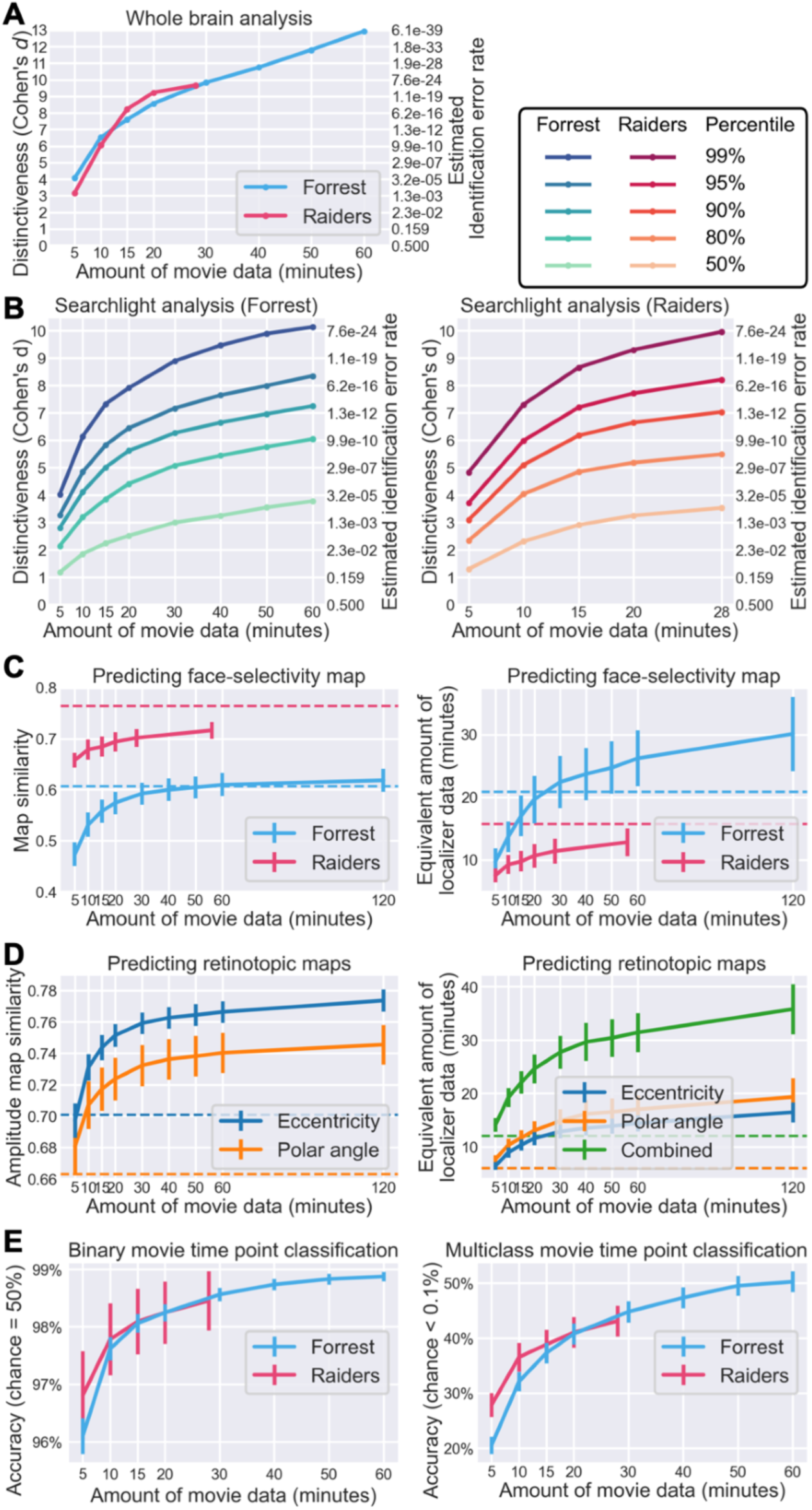
Effect of data volume on model performance. **(A)** Effect of data volume on the distinctiveness of an individual’s tuning matrix (cf. Figure 2D). With 10 minutes or more movie data, the within-subject similarity of tuning matrices were more than 6 standard deviations away from between-subject similarities on average, corresponding to a participant identification error rate of less than 1/10^9^. **(B)** Effect of data volume on the distinctiveness of local tuning matrices (cf. Figure 2E). Different lines denote different percentiles across searchlights, from an average searchlight (50^th^ percentile) to a highly distinctive searchlight (99^th^ percentile). **(C)** Predicting face-selectivity map with lower volumes of movie data (cf. Figure 3C). Face-selectivity maps can be accurately predicted with 20 minutes of movie data, but the prediction performance continues to grow with more data. Based on psychometrics and the quality of predicted maps, we estimated the amount of localizer data needed to achieve a similar quality (right panel). For the Forrest dataset, 30 minutes of movie data works better than standard localizers (21 minutes). Dashed horizontal lines indicate Cronbach’s alpha (left panel) or the actual duration of localizer scans (right panel). **(D)** Predicting retinotopic maps based on less movie data (cf. Figure 4C). **(E)** Quality of predicted response patterns for movie time points based on a model estimated from varying volumes of data (classification accuracy; cf. Figure 5C and 5D). Binary classification results on the left panel; multiclass results on the right panel. Both were based on 100 PCs. To summarize, the performance of our model continuously grows with more training data, but for certain tasks (e.g., individual identification, predicting category-selectivity and retinotopic maps), only a small amount of movie data (e.g., 30 minutes) is needed to achieve satisfying performance.

**Figure 7.**
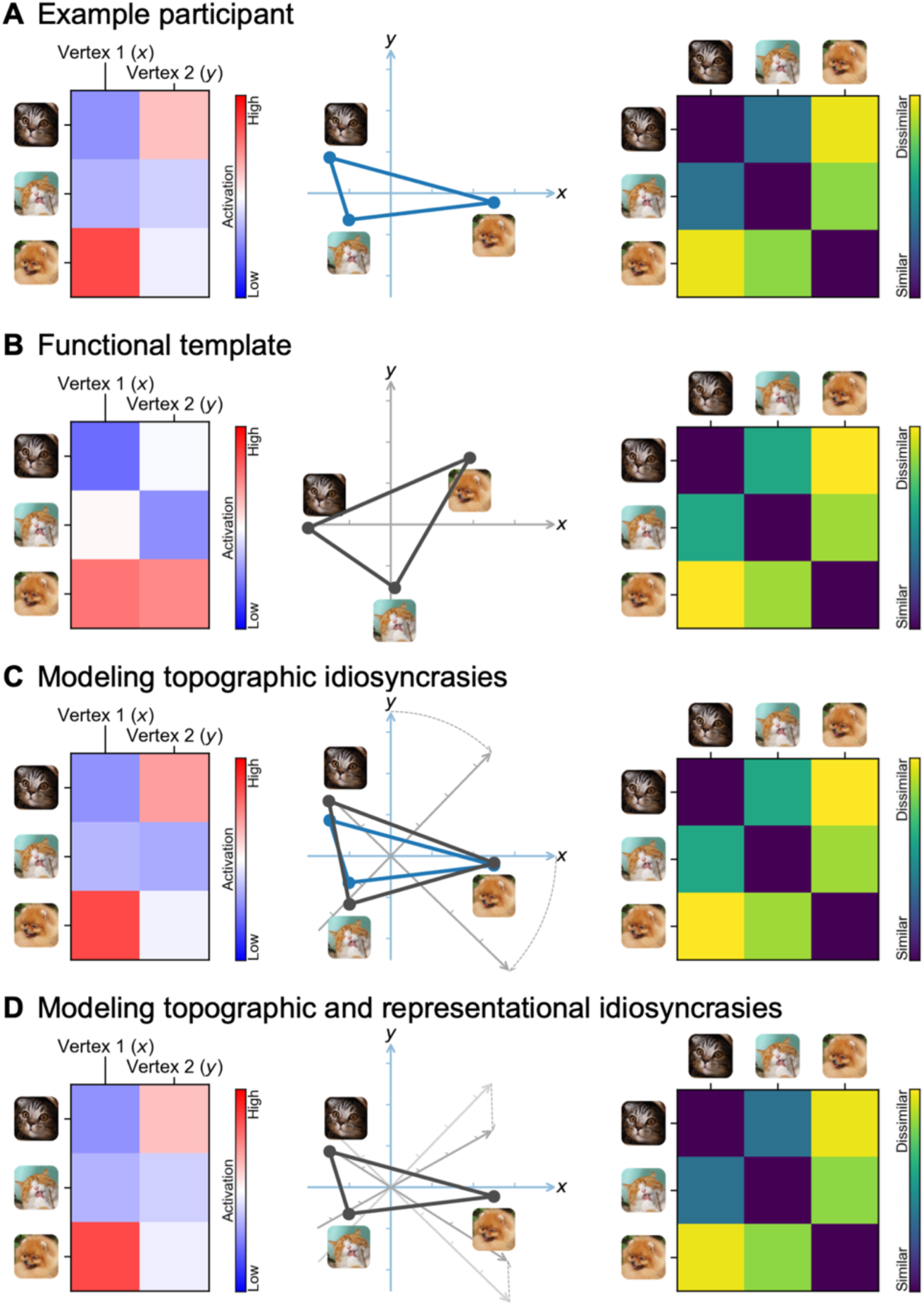
Schematic illustration of modeling a participant’s brain functional organization as a linearly transformed functional template. (**A**) A participant’s brain responses constitute a data matrix, where rows are stimuli (e.g., time points in a movie) and columns are cortical vertices (left). Multiple vertices form a high-dimensional space, where each vertex is a dimension, and each stimulus is a point in the space (middle). Information is encoded in the distances between the points. Such information can be summarized using a representational dissimilarity matrix (RDM), where each entry is the (dis)similarity between a pair of stimuli (right). (**B**) The RDM of the template resembles that of a participant (right), but the data matrix is usually quite different (left). This is because different brains encode the same information using different cortical topographies—the vertices collectively perform similar functions across individuals, but the function for each single vertex is quite different across individuals. (**C**) The participant’s idiosyncratic topographies can be predicted by a rotation of the template’s feature space (middle). The rotation changes the topographies of the template and makes the spatial patterns (rows of the data matrix) more similar to the participant’s (left), without changing the information content, or the RDM (right). (**D**) A linear transformation of the template can fully predict a participant’s responses by modeling both the participant’s idiosyncratic topographies and idiosyncratic information content; that is, both the “what” and “where” of a participant’s brain functional organization. Note that the schematic illustration is oversimplified; a typical fMRI data matrix contains thousands of stimuli/time points (rows) and tens of thousands of vertices (columns), and a real neural feature space is a high-dimensional space (hyperspace).

We observed a similar effect of data volume on functional distinctiveness in local brain areas based on a searchlight analysis (Figure 6B). The distinctiveness based on movie responses differs inherently across brain regions and is highest in temporal and occipital regions and lowest in somatosensory and motor regions (Figure 2E). Therefore, instead of a simple average value, we assessed key percentiles of the distribution. Specifically, we assessed the effect of data volume on the 50^th^, 80^th^, 90^th^, 95^th^, and 99^th^ percentiles of the distribution, representing local brain areas with low to high distinctiveness. With 15 minutes of movie data, the Cohen’s *d* for the 95^th^ percentile was 5.83 for the *Forrest* dataset and 7.19 for the *Raiders* dataset.

The prediction performance for face-selectivity maps also increases with more movie data (Figure 6C). For the *Forrest* dataset, the correlation between localizer-based and model-predicted maps was 0.557, 0.592, 0.610, and 0.618 for 15, 30, 60, and 120 minutes of movie data, respectively. For the *Raiders* dataset, the similarity was 0.684, 0.702, and 0.716 for 15, 28, and 56 minutes of data, respectively. Note that for the *Forrest* dataset, the similarity sometimes exceeded Cronbach’s alpha, which means the model-predicted map is more accurate than a map based on 4 localizer runs (21 minutes). The quality of localizer-based maps increases with more localizer data, which can be estimated using the Spearman–Brown prediction formula (Brown, 1910; Spearman, 1910). Based on Cronbach’s alpha and the Spearman–Brown prediction formula, we estimated the amount of localizer data needed to achieve similar accuracy as our model. For the *Forrest* dataset, the maps predicted by 15, 30, 60, and 120 minutes of movie data were as accurate as 17.0, 22.4, 26.2, and 30.1 minutes of localizer data, respectively. For the *Raiders* dataset, the maps predicted by 15, 28, and 56 minutes of movie data were as accurate as 9.7, 11.4, and 12.8 minutes of localizer data, respectively.

Note that brain responses to movies contain richer information than traditional experimental paradigms. Besides the face-selectivity map, many different maps can be estimated using the same movie data, such as retinotopic maps. With 15, 30, 60, and 120 minutes of *Forrest* data, the correlation between localizer-based and model-predicted amplitude maps were 0.744, 0.759, 0.766, and 0.774, respectively, for the eccentricity map; and 0.717, 0.732, 0.740, and 0.746, respectively, for the polar angle map (Figure 6D). These similarities were much higher than the corresponding Cronbach’s alpha values. Based on the Spearman–Brown prediction formula, the quality of the predicted maps was equivalent to 22.1, 27.7, 31.4, and 35.8 minutes of retinotopic scans, respectively.

The prediction performance for fine-grained response patterns to the movie also increases with the amount of movie data (Figure 6E). For the *Forrest* dataset, the accuracy for binary time point classification was 98.1%, 98.6%, and 98.9% for 15, 30, and 60 minutes of training movie data, respectively. For multiclass classification, the accuracy was 37.3%, 44.8%, and 50.3%, respectively. Similar results were observed for the *Raiders* dataset, where the binary classification accuracy was 98.1% and 98.5% for 15 and 28 minutes of training movie data, respectively, and the multiclass classification accuracy was 38.8% and 43.1%, respectively. To sum up, the performance of our model grows continuously with more data. For certain tasks (e.g., individual identification, predicting retinotopic maps), 10 to 20 minutes of movie data might be sufficient to achieve satisfying performance. Additional data will further improve the performance of our model, at least up to the typical duration of a feature film (2 hours).

## 3. Discussion

In this work, we present an individualized model of fine-grained brain functional organization. Through a series of analyses, we demonstrate that (a) the individualized tuning functions recovered by our model for each person are highly reliable across independent data; (b) our model can accurately predict an individual’s topographic brain responses to new stimuli, such as object categories and retinotopic localizers; (c) our model accurately predicts fine-grained response patterns to movies, which can be used to distinguish different time points (TRs) of the movie; and (d) the performance of our model continuously improves with more training data. Besides high reliability and high prediction accuracy, our model also shows high specificity—the predicted responses tuned to a given individual are much more similar to the actual responses for that person than predicted responses tuned to other individuals. To our knowledge, this is the first individualized model of brain function that offers vertex-level (voxel-level for volumetric data) spatial resolution. That is, our INT model provides out-of-sample generalization to new participants at the quality and spatial resolution of within-subject data acquisition.

Like most biological systems, the functional architecture of the brain is “degenerate”, such that roughly the same information can be instantiated in structurally different ways across different brains (Edelman and Gally, 2001; Haxby et al., 2020). In this work, we used searchlight hyperalignment algorithms (Guntupalli et al., 2016) to create a functional template of brain responses based on the training participants. The template is a common, high-dimensional response space, and its column vectors (response time series of features) span the space of response time series across vertices and participants. We took advantage of this property and created a set of basis vectors, so that we could express the response time series of each vertex and each participant as a linear combination of the same set of basis vectors. These weights offer a way to directly compare the functional architecture of different participants and different vertices. Based on these weights, we created the individualized tuning matrices that describe the brain functional organization of each participant, which can be used to accurately predict the participant’s idiosyncratic responses to various stimuli.

The present model provides a theoretical advance over previous hyperalignment algorithms by capturing not only topographic idiosyncrasies, but also inter-individual differences in representational geometry. The first component of the model introduces a new hyperalignment algorithm that we refer to as warp hyperalignment (WHA). WHA warps the representational geometry of one participant (or the template) to match the unique representational geometry of another participant, and thus it captures both topographic idiosyncrasies and representational idiosyncrasies. The second component of the model derives individualized tuning matrices in each participant’s native cortical topography from the WHA model, which we refer to as the Individualized Neural Tuning (INT) model. In contrast to our earlier hyperalignment algorithms for creating a common model information space with individual transformation matrices calculated using the Procrustes algorithm (which preserves representational geometry) (Busch et al., 2021; Feilong et al., 2021, 2018; Guntupalli et al., 2018, 2016; Haxby et al., 2020, 2001; Jiahui et al., 2020), WHA calculates transformations using ensemble regularized regression that allows for individualized representational geometries. WHA also introduces a new way to calculate a template matrix *M* in a single step that more accurately reflects the central tendency for cortical topography and is not biased towards the topography of a “reference brain”. The common model space in our previous models, *M*, had as many dimensions as cortical vertices (approximately 20,000 to 60,000). In the INT model, a change of basis from *M* to *S* recasts the common model space into a smaller orthogonal basis with approximately 3,000 dimensions. In our previous algorithms we studied individual differences in responses and connectivity as residuals around shared content in the model space, *M*. In the INT model, by contrast, we model individual differences in the transformation matrices, *T*, which capture individual differences in both content and cortical spatial topography of functional patterns in participants’ native cortical topographies. Because individual differences in representational geometry are now contained in the individual transformation matrices, *T*, the new model space, *S*, is a neural data-driven stimulus matrix that is not confounded with individual differences in representational geometry. Moreover, comparable estimates of *T* can be calculated from responses to different stimuli, giving the INT model more flexibility in its application, as well as greater precision. In our previous algorithms, we performed between-subject classification of response patterns after projecting all participants’ data into the common model space, *M*. In the INT model, we perform between-subject classification by comparing each test participant’s response pattern in their native space to response patterns from other participants projected into that test participant’s native space.

The INT model separates neural responses into stimulus-related information and stimulus-general neural tuning, which can be estimated separately. The stimulus-related information is represented as the stimulus matrix *S*, which is derived based on the neural responses of the training participants when the functional template is created. After the functional template has been created, descriptors for additional stimuli can be estimated based on a subset of training participants for whom responses to the new stimuli are available. These descriptors for new stimuli extend the original stimulus matrix *S*, and they can be used to predict individualized responses to the new stimuli for the left-out test participants. For example, we built the functional template based on responses to the movie, estimated the stimulus descriptors for object categories and retinotopic localizers, and used these stimulus descriptors to estimate the category selectivity maps and retinotopic maps of left-out test participants. In other words, the original stimulus matrix *S* can be extended based on a subset of participants, provided that we have their neural responses to the new stimuli and their tuning matrices. On the other hand, the tuning matrix *T* of a new participant, which represents stimulus-general neural tuning, can be accurately estimated with several minutes of movie data (Figure 6). Therefore, our INT model makes it possible to accurately predict individualized, out-of-sample responses to a wide range of stimuli based on a rich normative functional template and a relatively small amount of fMRI data from a new participant.

A major objective of studying individual differences in brain functional organization is to build biomarkers that are associated with cognition, behavior, and disorders. Our model focuses on semi-shared components of brain functional organization and is ideal for this purpose. By “semi-shared” we mean that the same component exists in multiple brains but differs in amplitude and topography. These reliable variations across individuals may covary with phenotypes of interest and provide accurate biomarkers. A fully shared component, which is identical across brains, cannot covary with other variables by definition. A fully idiosyncratic component that only exists in one brain, on the other hand, cannot be used to build generalizable models. For example, a specific component that only exists in one schizophrenic brain may be of interest for a case study but cannot be used to diagnose other schizophrenic individuals because it doesn’t exist in other brains. Our model focuses on how the same set of components are instantiated in different forms across the functional organization of different brains. Given the large number of components (over 3,000 in the current implementation) and observation that they vary across brains in a variety of ways, these semi-shared components provide a promising basis for developing biomarkers. Similar to our previous work (Feilong et al., 2018), brain regions that have the most shared and synchronized responses (Guntupalli et al., 2016; Hasson et al., 2010, 2004) are also the regions showing the most reliable differences, suggesting the great potential of using semi-shared components to study individual differences.

In this work we evaluated our model using two different movie datasets, both of which yielded highly similar results. The *Forrest* dataset was collected using a 3 T Philips Achieva dStream MRI scanner in Germany, with German-language audio, a TR of 2 seconds, and a spatial resolution of 3 mm. The *Raiders* dataset was collected using a 3 T Siemens Magnetom Prisma MRI scanner in the US, with English-language audio, a simultaneous multi-slice acceleration factor of 4, a TR of 1 second, and a spatial resolution of 2.5 mm. Despite these differences, our model worked well for both datasets, suggesting it is robust over differences in scan parameters and other details. Recently many large-scale neuroimaging datasets have become openly available (Alexander et al., 2017; Horien et al., 2020; Nastase et al., 2021; Snoek et al., 2021; Taylor et al., 2017), and many have naturalistic movie-viewing sessions similar to our datasets. The synergy between our individualized model of brain function and large-scale neuroimaging datasets offers a great opportunity to study individual differences in brain functional organization and their correlates with various phenotypes.

In this work we focused on neural response profiles to the movie. However, in theory, the algorithm itself can be applied to any kind of data matrices. In our previous hyperalignment algorithms, the searchlight procedure originally developed based on response profiles (RHA) (Feilong et al., 2018; Guntupalli et al., 2016; Haxby et al., 2020; Jiahui et al., 2020) has been applied successfully to connectivity profiles (CHA) (Feilong et al., 2021; Guntupalli et al., 2018; Nastase et al., 2020) and a hybrid of both (H2A) (Busch et al., 2021); and the original algorithm developed based on fMRI data of humans (Haxby et al., 2011) has been applied successfully to electrophysiology recording data of rodent neurons (Chen et al., 2021). We leave it to future works to assess the generalizability of the INT model to other functional profiles, modalities, and species.

## 4. Methods

### 4.1. Overview of the INT model

The fine-grained functional architecture of the brain encodes rich information (Haxby et al., 2020, 2014, 2001) and affords reliable measures of individual differences in brain functional organization that are predictive of differences in behavior (Feilong et al., 2021, 2018). In this work, we present the individualized neural tuning (INT) model, an individualized model of fine-grained brain functional organization, to better model these differences. The INT model has three key features. First, it has fine spatial granularity, which affords access to the rich information encoded in vertex-by-vertex (or voxel-by-voxel) patterns. Second, it models each individual’s idiosyncratic functional organization as well as that individual’s topographic projection onto the cortex, and thus it can be used to study both functional differences and topographic differences. Third, it models the individualized response tuning of cortical vertices in a way that generalizes across stimuli, and therefore the model parameters can be estimated from different stimuli, such as different parts of a movie that have different durations. These three features make the INT model a powerful tool to study individual differences in fine-grained functional organization of the brain.

The INT model is based on the conceptual framework of hyperalignment (Guntupalli et al., 2018, 2016; Haxby et al., 2020, 2011). Hyperalignment models the fine-grained functional organization of each brain as a high-dimensional feature space, and it creates a high-dimensional common space based on the shared functional profiles of a group of participants. Hyperalignment also provides a way to transform between different spaces using a high-dimensional rotation, which can be used to project the data from the common space to a participant’s native anatomical space, from a participant’s space to the common space, or from a participant’s space to another’s (Jiahui et al., 2020). This high-dimensional rotation resolves topographic differences, which is critical to study individual differences in fine-grained functional organizations (Feilong et al., 2021, 2018).

The INT model starts with creating a functional template *M* (a matrix of shape *t* × *v*) based on the data of the training participants (*n –* 1 for leave-one-subject-out cross-validation), which corresponds to the hyperalignment common space. The template *M* has the same shape as the data matrix *B* of a participant, and its function and topographies are representative of the group of participants used to create the template. The data matrix *B_(p)_* of the participant *p* is modeled as a matrix multiplication of the shared functional template *M* and an idiosyncratic linear transformation *W_(p)_* (*v* × *v*). We use a new hyperalignment algorithm (“warp hyperalignment”, WHA) to derive the transformation instead of Procrustes-based hyperalignment, so that the transformation is a linear transformation instead of an improper rotation. An improper rotation (rotation and reflection) changes how the information is encoded on the cortex (“where”) but it does not change the content information (“what”), and thus it only accounts for topographic differences across individuals. A linear transformation allows scaling and shearing, which also warp the representational geometry of the template to model the idiosyncratic representational geometry of each participant, and therefore it accounts for both topographic (“where”) and functional (“what”) differences.

With warp hyperalignment, we obtain a modeled data matrix 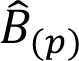, which are the brain responses that can be accounted for by the functional template and the linear transformation (i.e., *MW_(p)_)*. To derive a measure of neural response tuning that generalizes across stimuli, we decompose 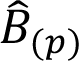 into two matrices, a stimulus matrix *S* (*t* × *k*) shared by all participants, and a tuning matrix *T_(p)_* (*k* × *v*) that is specific to the participant *p*. With the decomposition, the temporal information, such as contents of a movie over time, is factored into *S*. In the tuning matrix *T_(p)_*, the response tuning function of each cortical vertex is depicted using a column vector of *k* elements, which is the same for all stimuli.

To sum up, with the INT model we use the tuning matrix *T_(p)_* to model each participant’s individualized functional organization. The tuning matrix has a fine-grained spatial granularity, models the participant’s topographic and functional idiosyncrasies, and generalizes across stimuli. In the next few sections, we describe in detail the steps we used to derive the tuning matrices and to benchmark the reliability, validity, accuracy, and specificity of our INT model.

### 4.2. Building the functional template

In each cross-validation fold, we built a functional template based on the training participants and modeled each test participant’s data matrix as the linearly transformed template in a high-dimensional space. Both the data matrix and the functional template have the same shape *t* × *v*; that is, the number of time points by the number of cortical vertices. The template was created in a way that its functional properties—both in terms of representational geometry and cortical topography—are representative of the training participants.

#### 4.2.1. Searchlight-based algorithm

We built the template using a searchlight-based algorithm. For each searchlight, we built a local template based on all vertices within the searchlight. We then combined all the local templates into a whole brain template. Each local template contains modeled response profiles of vertices in the corresponding searchlight. Each vertex is included in multiple searchlights, and each searchlight and the corresponding local template offers a modeled response profile for the vertex. We combined these modeled response profiles of the same vertex into a single response profile for the vertex, which is the vertex’s response profile in the whole brain template. In our previous algorithms, we combined local searchlight templates by adding together the modeled response profiles of the vertex to form the final response profile of the vertex (Guntupalli et al., 2018, 2016). In this work, we instead used a distance-based weighted average instead of summation. Specifically, the weight was computed as 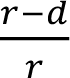, where *r* is the searchlight radius (20 mm), and *d* (0 <= *d* <= *r*) is the distance between the vertex and the center of the searchlight. In other words, the weight is 1 when the center of the searchlight is the vertex itself, and close to 0 when the vertex is close to the boundary of the searchlight. This improved procedure makes the searchlights closer to the vertex contribute more to the final modeled response profile of the vertex (due to weighting local templates), and the scale of the modeled response profile for a vertex similar to the actual response profile for that vertex (due to using averaging instead of summation).

#### 4.2.2. Building local templates

In order to estimate the INT model, we must first create a functional template capturing the consensus functional organization (which we refer to as *M*). Within each searchlight, we created a local template using a PCA-Procrustes algorithm, and the matrix shape of the local template is the same as a local data matrix (i.e., the number of features is the same as the number of vertices in the searchlight, not the total number of vertices). First, we concatenated all training participants’ data matrices in the searchlight along the features dimension to form a group *n* × *v;* that is, the number of participants times the number of vertices in the searchlight. We then applied principal component analysis (PCA) to this concatenated data matrix. To keep the total variance the same for a single participant’s local data matrix and the local template, we divide the PC time series by √𝑛. Similar to our previous work (Haxby et al., 2011), here we chose to make the dimensionality of the local template the same as a single participant’s local data matrix, thus retaining the first *v* PCs and discarding the remaining. Note that the PCA is based on the data of all training participants, and thus the PCs summarize across all vertices and participants; each PC is a weighted sum of all vertices (in a given searchlight) across all training participants. The PCs capture the representational geometry for a given in searchlight in a way that is representative of the representational geometries of the training participants. In other words, the PCs provide a template that models the shared function of the searchlight.

We then used the orthogonal Procrustes algorithm to “align” the PCs to the training participants’ data, so that the functional topographies of the local template are also representative of the training participants. Mathematically, we want to find a rotation matrix *R* which minimizes the topographic differences without changing the information content.

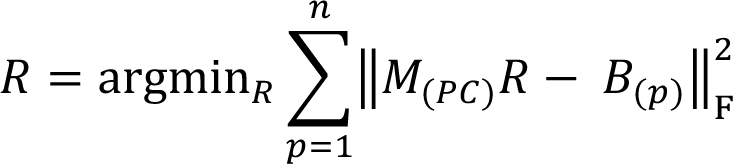

In this equation 𝑀_(*PC*)_ is the PC matrix, *B_(p)_* is the local data matrix of the *p*-th participant, *n* is the number of participants, and ‖ ‖_F*_ is the Frobenius norm.

To find the solution *R*, we applied the orthogonal Procrustes algorithm to concatenated data matrices. This time, we concatenated all training participants’ data along the samples (i.e., time points) dimension to form another group data matrix, where the number of rows is *n* × *t;* that is, the number of participants times the number of time points. We copied the template PC matrix *n* times and concatenated them in the same way, so that the concatenated PC matrix had the same shape as the concatenated group data matrix. We applied the orthogonal Procrustes algorithm to these two data matrices to get a rotation matrix *R*.

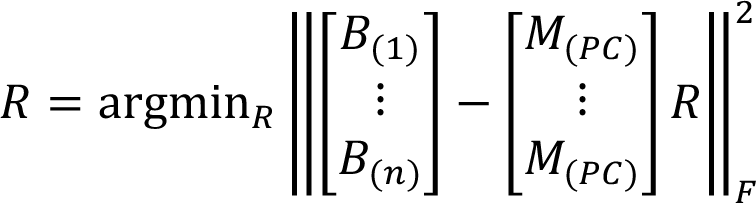

Note that the solution for this formula is the same as the previous one. However, because the matrices have been concatenated, the solution of the orthogonal Procrustes algorithm can be computed directly based on the singular value decomposition of the covariance matrix, which provides an analytical solution to the problem.

Similar to Procrustes-based hyperalignment algorithms, this rotation matrix *R* does not change the representational geometry or the information content in the data matrix. Instead, it changes the functional topographies so that one data matrix is “aligned” to another. In this case, a single rotation is estimated that best aligns the coordinate axes (i.e., PCs) of the template matrix and the coordinate axes (i.e., cortical vertices) of all participants, so that the functional topographies of the rotated template matrix maximally resemble those of the training participants. The final local template *M* is the PC matrix multiplied by the rotation matrix *R*: 𝑀 = 𝑀_(*PC*)_𝑅.

In short, we used the PCA-Procrustes algorithm to create a local template for each searchlight, which is representative of the training participants both in terms of representational geometry and cortical topography. The PCA step ensures that the functional profiles and representational geometry of the local template are close to those of the training participants, and the orthogonal Procrustes step ensures that the topographical distribution of these functions on the cortex is also representative of the training participants. After iterating over all searchlights, the local templates were combined into a single whole brain template using the distance-based weighted average method described above.

### 4.3. Modeling response tuning functions

We modeled each participant’s response data matrix *B_(p)_* as the template data matrix *M* multiplied by a linear transformation *W_(p)_*, plus some noise *E*:

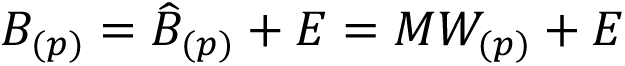

Unlike Procrustes-based hyperalignment (Haxby et al., 2011), in which the transformation matrix *W_(p)_* (often denoted as *R)* is a rigid improper rotation, the linear transformation *W* allows warping of representational geometry. Consequently, individual differences in representational geometry are embedded in the transformation matrices, *W*, rather than in the individual information projected into the model space, *M*. We name the new algorithm “warp hyperalignment” (WHA) to emphasize its capacity to warp representational geometries and to distinguish it from previous algorithms.

We computed the linear transformation *W_(p)_* using a searchlight-based algorithm, similar to the procedure we used to create the template *M*. That is, for each of the searchlights, we computed a local transformation, and these local transformations were combined using the distance-based weighted average.

Typically, a model needs to be regularized to avoid overfitting and to increase its generalizability to new data. For the orthogonal Procrustes algorithm, the linear transformation *W_(p)_* is constrained to be orthogonal (i.e., an improper rotation in a high-dimensional space), which can be considered as a strong regularization. In this work, we allowed scaling and shearing in the transformation, which models individual differences in function, such as representational geometry. We used two methods to avoid overfitting in model estimation. First, we used ridge regression with a regularization parameter of 10^3^ based on independent pilot data not presented here. Second, we used an ensemble method which we call k-fold bagging. That is, for each participant and each searchlight, we trained 100 ridge regression models based on bootstrapped samples (bootstrapped time points; sampled with replacement), and we averaged the weights of these 100 models to serve as the weights for the final model (described in detail below).

#### 4.3.1. Ensemble ridge regression models

We used ensemble learning (Zhou, 2012) to improve the accuracy and generalizability of our models. Specifically, we adapted the bootstrap aggregating (“bagging”) algorithm (Breiman, 1996) for our time series data. Bagging is commonly used to reduce model variance and avoid overfitting by averaging across models trained on bootstrapped samples. It also provides estimation of model performance on new data through out-of-bag cross-validation. During out-of-bag cross-validation, the predicted value of a data point is the average prediction of models that were not trained with the time point (i.e., out-of-bag models). In this case, this data point serves as the test data and the other time points as training data. Typically, bootstrapped samples are randomly drawn with replacement from the original sample. A participant’s fMRI data (e.g., responses to movies) usually comprises hundreds or thousands of time points. With the classic bagging algorithm, it often happens that some time points are drawn by all bootstrapped samples, which makes them inappropriate for model evaluation using out-of-bag cross-validation (i.e., no out-of-bag models for these data points). To use as much data as possible for cross-validation, we augmented the classic bagging algorithm with a k-fold scheme.

In each k-fold repetition, we first divide all time points randomly into k folds. For a given fold, we set aside the data in that fold to serve as candidate test data, while data in the other k – 1 folds serve as candidate training data. We then drew a bootstrapped sample from the candidate training data and used it to train a model. This procedure guarantees that the candidate test data can be used for model evaluation because they were not used in model training. Some candidate training data may not get chosen by the bootstrapped sample and these data also serve as test data for model evaluation. In other words, for each model, the actual test data includes both candidate test data and the candidate training data not drawn by the bootstrapped sample. After an iteration over all k folds, we obtained k trained models. For each data point, our resampling procedure ensures that at least one of the k models was not trained with the data point. In this work, we used k = 5 and repeated the k-fold scheme for 20 times, and thus the prediction for each data point was the average of at least 20 out-of-bag models.

To account for temporal autocorrelation caused by the hemodynamic response function, we also introduced temporal “buffers” for out-of-bag cross-validation. That is, when we evaluate model performance on a certain time point, we exclude not only models trained with the time point itself, but also models trained with time points less than 10 s away from the time point used for evaluation. For example, for a 2 s TR length, when we evaluate model performance for the *i*-th TR, we exclude models trained with any of the 11 TRs from *i* – 5 to *i* + 5. To avoid removing too many buffer time points from the training data, we divided time points into groups by grouping them into 10 s segments (5-TR segments for a 2 s TR), and assigned all time points in the same segment to the same fold.

The adapted bagging algorithm and the out-of-bag cross-validation procedure were only based on the training data (for the test participant). Similar to the inner-loop of nested cross-validation, the training and test folds discussed in this context were both part of the training data. Because independent data were used in out-of-bag evaluation, this procedure provides an unbiased way to estimate model performance on new data, such as the actual test data.

#### 4.3.2. Separating stimulus and tuning information

Based on the whole-brain functional template *M* and the linear transformation *W_(p)_* derived by warp hyperalignment, we obtained a modeled brain response matrix 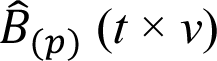 for the participant *p*, which are the responses of the participant that can be accounted for by the linearly transformed template. To model the participant’s neural response tuning independent of stimulus information, we derived a tuning matrix *T_(p)_* (*k* × *v*) by a matrix decomposition of 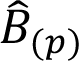. This matrix decomposition factors the temporal information into the matrix *S* (*t* × *k*). The columns of *S* are a set of basis response profiles (i.e., response time series to the movie). The response profile of each vertex is modeled as a linear combination (i.e., weighted sum) of the basis profiles, and the weights of the linear combination are the corresponding column in *T_(p)_*, which is a column vector of *k* elements. This column vector is independent of the stimulus, and it reflects the response tuning function of the vertex. We refer to this column vector as the tuning profile of the cortical vertex to distinguish it from the response profile (response time series). To use the tuning matrices to model differences in neural tuning across vertices and across individuals, ideally the tuning matrices should have several properties: (a) cortical vertices that have larger differences in response time series also have larger differences in their tuning profiles; (b) individuals who are more similar based on their response profiles are also more similar based on their tuning matrices; (c) the same tuning matrix can be estimated from different stimuli, such as different parts of the movie with different durations. These objectives motivate us to find a matrix *S* with three properties: (a) the columns are orthogonal to each other; (b) each column has unit variance; and (c) the columns of *S* form a basis set of response profiles. Orthogonality is necessary to make *S* a similarity transformation, so that differences in *T_(p)_* across vertices and across individuals are proportional to their differences in 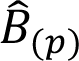. Unit variance ensures that the scale of the estimated *T_(p)_* is the same for different amounts of data, such as data matrices from different parts of the movie. That the columns of *S* form a set of basis response profiles means the response profile of each vertex and each participant can be expressed as a linear combination of the basis profiles. In other words, *S* can be used to fully model *B_(p)_* and 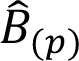 without any loss of information.

There are many choices of *S* which have all these properties and work similarly well for our purposes. In this work, we use the normalized principal components (PCs) from a group-PCA. The normalized PCs work well in practice, as is shown by the benchmarking analyses. Furthermore, due to the nature of PCA, they provide an easy way to reduce data dimensionality when less dimensions are desired. In this work we did not reduce dimensionality, and thus *k* equals the rank of the concatenated matrix, which is the same as the number of time points in the movie in practice (approximately 3000). We performed the group-PCA using a singular value decomposition (SVD) on the concatenated data matrices of all participants, and rescaled the first matrix *U* to get *S*.

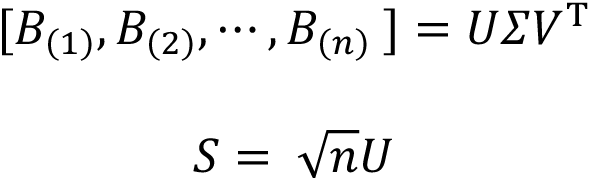

Based on the conceptual framework of hyperalignment (Haxby et al., 2020, 2011), different brains share the same functional basis. In practice, the shared functional basis is instantiated as a hyperalignment common space, which is a functional template. The response profiles of the template’s vertices form a set of basis response profiles, and the response profile of each cortical vertex is expressed as a linear combination of these basis response profiles. The weights of the linear combination are the elements in the corresponding column of the transformation matrix. Note that the transformation matrix based on the searchlight algorithm is highly sparse, and the weights of the linear combination are non-zero only for local neighborhoods of vertices (i.e., vertices included in the same searchlight) in the template. As a result, the response profile of each vertex is modeled using a different set of vertices, whose response profiles highly covary due to spatial autocorrelations.

In the INT model, the columns of matrix *S* serve as the set of basis response profiles, which are orthogonal vectors with unit variance. The response profiles of all vertices and all participants are all expressed as a linear combination of the same basis set, which affords the study of functional tuning differences across vertices and across individuals based on tuning matrices, whose columns comprise the linear combination weights. In other words, we are replacing local basis sets (response profiles of adjacent vertices) with a single global basis set of response profiles (columns of *S*). Conceptually *S* is also a common space, but different from *M*, the features in *S* are completely virtual and do not correspond to specific cortical loci.

The features in *S* are neural data-driven stimulus descriptors. They are derived from shared brain responses and reflect the primary ways cortical vertices response to stimuli. Each stimulus (e.g., movie time point) is described as a row in *S*, which is a vector of *k* elements, and each element indexes to what extent a virtual feature responds to the stimulus. In other words, the row vector describes the key features of the corresponding stimulus based on neural responses. Therefore, here and elsewhere we refer to *S* as the stimulus matrix.

Because stimulus information is factored into *S*, the information in the tuning matrix *T_(p)_* is neural response tuning of cortical vertices that is the same for a wide variety of stimuli from the space spanned by a naturalistic, audiovisual movie stimulus. For example, when we divide the neural response data matrix *B* into two halves, each half can be modeled using the corresponding half of *S* and the same *T_(p)_* (Figure 1B). This property has an important implication for *T_(p)_*: Once the functional template is created, the same individualized *T_(p)_* can be estimated from independent data of the same individual (e.g., different parts of a movie), and the amount of data used to estimate *T_(p)_* can be less than the amount of data used to create the functional template (e.g., responses to part of the movie instead of the entire movie).

Furthermore, the INT model can be extended to model responses to stimuli that were not used to create the template. Given the neural responses to new stimuli from a group of participants (which can be a subset of all participants) and their tuning matrices, the stimulus descriptors *S_(new)_* for the new stimuli can be estimated (Figure 1C) and used to predict other participants’ responses to the new stimuli.

In the sections below, we use a series of analyses to demonstrate the reliability, validity, accuracy, and specificity of our INT model. In the first analysis, we show that the tuning matrices estimated from different parts of the movie are highly similar for the same individual but dissimilar for different individuals. In the second and third analysis, we show that individualized responses to new stimuli (category selectivity and retinotopic maps) can be accurately predicted by estimating the stimulus descriptors for the new stimuli. In the fourth analysis, we show that the INT model can accurately predict individualized fine-grained spatial response patterns, such as responses to a specific time point of a movie. In the fifth analysis, we show that 10–20 minutes of movie data are sufficient for satisfying performance of the INT model, but the performance grows continuously with more data.

### 4.4. Datasets

#### 4.4.1. The Forrest dataset

The *Forrest* dataset is part of the Phase 2 data of the studyforrest project (Hanke et al., 2014). It contains 3 T fMRI data collected from 15 right-handed German adults (mean age 29.4 years, 6 females) during movie watching, retinotopic mapping, and object category localizers (Hanke et al., 2016; Sengupta et al., 2016). Each participant’s movie data comprised eight runs of approximately 15 minutes each, while the participant watched a shortened version of the audiovisual feature movie *Forrest Gump*. In total, 3599 volumes were collected over the course of 2 hours of scanning. The retinotopic data comprises four 3-minute runs (12 minutes in total), and the four runs corresponded to expanding rings, contracting rings, clockwise wedges, and counterclockwise wedges. The object category localizer data contains 4 runs that are 5.2 minutes each (20.8 minutes in total). Each run contains two 16 s blocks for each of the 6 categories (bodies, faces, houses, objects, scenes, and phase scrambled images). During each block, 16 grayscale images were displayed for 900 ms each with a 100 ms interval. During the object category localizer scans, the participant performed a central letter reading task to maintain attention and fixation. All these data were acquired with a Philips Achieva dStream MRI scanner and a gradient-echo EPI sequence, with which a whole brain image containing 3 mm isotropic voxels was acquired every 2 seconds. More details of these datasets can be found in the data descriptors for the 3 T studyforrest data (Hanke et al., 2016; Sengupta et al., 2016).

#### 4.4.2. The Raiders dataset

The *Raiders* dataset contains data from 23 participants (mean age ± SD: 27.3 ± 2.4 years; 12 females) while they were watching the second half of the movie *Raiders of the Lost Ark* (Nastase, 2018). The movie scan comprised 4 runs that were 14–15 minutes each (850, 860, 860, and 850 seconds, respectively). In total, 3420 volumes were collected for each participant, with a 1 second TR and 2.5 mm isotropic voxels. The movie clips of adjacent runs had 20 seconds of overlapping content, and thus we removed 10 seconds of data from the end of first run and 10 from the beginning of the second run during analysis. After chopping off the overlapping content, the remaining movie data were 14 minutes (840 TRs) per run and 56 minutes in total. Among the 23 participants, 20 also had localizer data. The localizer data were the same data used in (Jiahui et al., 2020). It was collected using the same scan protocol as the movie, and it comprised four runs of 3.9 minutes each (15.6 minutes in total). Each run comprised 10 blocks, 2 per category (faces, bodies, scenes, objects, and scrambled objects), and each block was 18 seconds long. Each block comprised 6 video clips that were 3 seconds each. During the localizer scans, the participant performed a 1-back repetition detection task based on the video clips. The *Raiders* dataset was collected using a 3 T Siemens Magnetom Prisma MRI scanner with a 32-channel head coil at the Dartmouth Brain Imaging Center, with the same scan protocols as (Visconti di Oleggio Castello et al., 2020). Each second, a volume was collected with 2.5 mm isotropic voxels and whole brain coverage. The volume comprised 52 axial slices collected in an interleaved fashion with gradient-echo echo-planar imaging. Each slice had a 96 × 96 matrix and an FOV of 240 × 240 mm^3^. The TE was 33 ms, flip angle was 59°, and the phase encoding direction was anterior–posterior. The imaging was accelerated using a simultaneous multi-slice (SMS) factor of 4 and no in-plane acceleration. All participants gave written, informed consent, and were paid for their participation. The study was approved by the Institutional Review Board of Dartmouth College.

#### 4.4.3. MRI Preprocessing

We ran fMRIPrep (Esteban et al., 2019) on all MRI data, using version 20.1.1 for the *Forrest* dataset, and 20.2.0 for the *Raiders* dataset. After fMRIPrep, functional data from all participants were projected onto a cortical surface and were in alignment with the fsaverage template (Fischl et al., 1999) based on cortical folding patterns. We then performed downsampling and nuisance regression in the same way as (Feilong et al., 2018). First, we downsampled functional data to a standard cortical surface mesh with 9372 vertices for the left hemisphere and 9370 vertices for the right hemisphere (approximately 3 mm vertex spacing; 10242 per hemisphere before removing non-cortical vertices). Then, we performed a linear regression to partial out nuisance variables from functional data separately for each run. The nuisance regressors include 6 motion parameters and their derivatives, global signal, framewise displacement (Power et al., 2014), 6 principal components from cerebrospinal fluid and white matter (Behzadi et al., 2007), and polynomial trends up to the 2^nd^ order. Finally, we normalized the residual time series of each vertex to zero mean and unit variance.

### 4.5. Assessing the reliability and specificity of tuning matrices

To make the tuning matrices a useful measure of brain functional organization, they need to have high reliability and specificity. That is, tuning matrices of the same individual based on independent data should be similar, and tuning matrices from different individuals should be dissimilar. Therefore, we split each participant’s movie data into two parts, and estimated a tuning matrix based on each part of the movie.

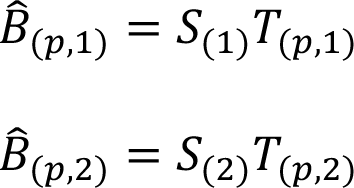

Where 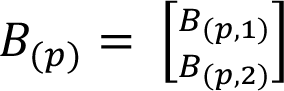, and 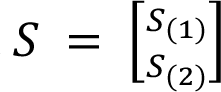. *T_(p,_*_1)_ and *T_(p,_*_2)_ are both estimations of *T_(p)_*, but they are estimated based on different parts of the movie (independent data).

To assess the reliability and specificity of the modeled tuning matrices, we computed a cross-movie-part similarity matrix for each dataset based on the estimated tuning matrices. The matrix has a shape of *n* × *n*, where each row corresponds to a tuning matrix based on the first part of the movie, each column corresponds to a tuning matrix based on the second part of the movie, and each entry is the correlation-based similarity between the two matrices. The diagonal of the matrix is the within-subject similarities, and the off-diagonal elements are between-subject similarities. A clear difference between diagonal and off-diagonal elements indicates a substantial difference between within-subject and between-subject similarities.

#### 4.5.1. Multi-dimensional scaling

To better visualize the similarities between estimates tuning matrices, we performed multi-dimensional scaling (MDS) using the T-distributed Stochastic Neighbor Embedding (t-SNE) algorithm (Van der Maaten and Hinton, 2008). We used a full individual differences matrix (i.e., 2*n* × 2*n* elements, comprising both same-movie-part and cross-movie-part dissimilarities based on correlation distance) as input to the t-SNE algorithm. The 2*n* tuning matrices were projected to a 2D space by t-SNE. Given any MDS algorithm would unavoidably distort distances during the projection, we used a perplexity parameter of 10 to reduce the distortions of distances between closer neighbors, which in this case are within-subject dissimilarities and several smallest between-subject dissimilarities. These dissimilarities are key to determine whether an individual can be easily identified based on the tuning matrix and a nearest-neighbor classifier.

#### 4.5.2. Distribution of tuning matrix similarities

For each tuning matrix, we extracted its within-subject similarity and between-subject similarities based on the cross-movie-part similarity matrix. These similarities correspond to the diagonal (within-subject) and off-diagonal (between-subject) elements of a row of the similarity matrix. We plotted the distribution of the within-subject similarity and between-subject similarities for each tuning matrix in Figure 2C, sorted by within-subject similarity.

#### 4.5.3. Distinctiveness index

For all tuning matrices, we found that within-subject similarity was far greater than the distribution of between-subject similarities. In other words, any participant can be identified by the modeled tuning matrix with an accuracy of 100% based on a simple one-nearest-neighbor classifier. To better describe how distinctive an individual is based on the modeled tuning matrix, we computed the distinctiveness index based on Cohen’s *d*:

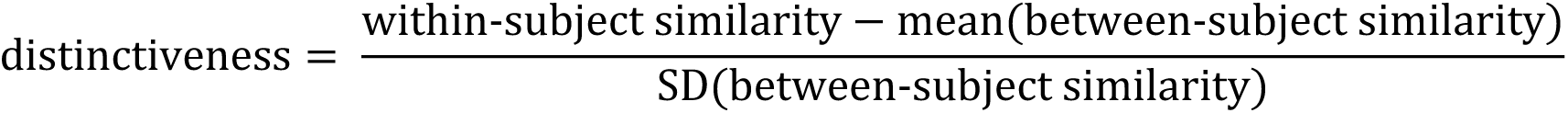

The distinctiveness index is a measure of effect size, and thus is comparable across datasets with different sample sizes. The similarities used to compute the distinctiveness index were Fisher-transformed correlation similarities, and therefore they approximately follow a normal distribution, and the distinctiveness index can serve as a *z*-statistic. Using the cumulative distribution function of the standard normal distribution, an identification error rate can be estimated based on the distinctiveness index.

#### 4.5.4. Searchlight analysis

To locate the brain regions where the functional organization is most distinctive, we performed a searchlight analysis (Kriegeskorte et al., 2006) using a searchlight radius of 20 mm. Within each searchlight, we computed a distinctiveness index for each tuning matrix based on vertices in the searchlight, and we averaged the distinctiveness index across all tuning matrices to get an average distinctiveness index for the searchlight. We repeated this process for each searchlight and obtained an average index for each searchlight. These average distinctiveness indices formed a map of distinctiveness for each dataset (Figure 2E).

### 4.6. Predicting category-selectivity maps

The previous analyses have shown that our model has high reliability and specificity. The modeled brain functional organization is highly similar for the same individual (based on independent data), and much less similar for different individuals. In this part, we tested the generalizability of our model. Specifically, we tested whether our model could predict responses to new stimuli that were not used in model training. Therefore, we trained our model based on the movie data and tested whether the model can be used to predict responses to various object categories. Here we use the “faces” category as an example to illustrate the procedure of our analysis, and the same procedure was applied to other object categories.

#### 4.6.1. Quality of localizer-based maps

The *Forrest* dataset has 4 static object category localizer runs per participant (for all participants), and the *Raiders* dataset has 4 dynamic object category localizer runs per participant (for 20 out of the 23 participants). For each run of each participant, we used general linear model to estimate the contrast of interest (faces vs. all other categories) and obtained a map of *t*-statistics for the contrast. Due to the presence of noise in localizer data, the estimated face-selectivity map is a combination of a “true” face-selectivity map of the participant and some noise. The component from the “true” map is supposed to be shared by all localizer runs, and thus the data quality and the level of noise can be estimated based on the similarity between the 4 maps (i.e., one from each run). We used Cronbach’s alpha to estimate the quality of the average map of the 4 runs. If we were to collect another 4 localizer runs from the participant and get a new average map based on the 4 new runs (i.e., independent data), then the expected correlation between the two average maps would be Cronbach’s alpha. In other words, if the correlation between the model-predicted map and the localizer-based (average) map is higher than Cronbach’s alpha, then the model-predicted map is more accurate than the average map based on 4 runs.

#### 4.6.2. Model-predicted category selectivity maps

We used a leave-one-subject-out cross-validation scheme to evaluate model performance. We built the template based on the *n* – 1 training participants’ movie data. We then computed a tuning matrix *T_(p)_* for each of the *n* participants based on the movie data. We modeled the face selectivity map as the brain response pattern to the specific “faces” category:

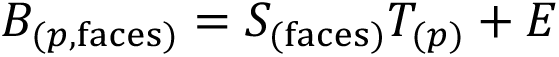

Here 𝐵_(*p*,faces)_ denotes the face-selectivity map for participant *p*, and 𝑆_(faces)_ denotes the stimulus descriptors for the “faces” versus other categories contrast. In this case, both 𝐵_(*p*,faces)_ and 𝑆_(faces)_are row vectors because there is only one stimulus (category). Both 𝐵_(*p*,faces)_ and *T_(p)_* were known for the training participants, and thus 𝑆_(faces)_ can be estimated using a general linear model (e.g., ordinary least squares) by finding the 𝑆_(faces)_ that minimizes the Frobenius norm. ∥𝐵_(*p*,faces)_ − 𝑆_(faces)_𝑇_(*p*)_∥_F_. This solution can be computed using ordinary least squares (“vanilla” regression), but here we used ensemble linear ridge regression to increase the accuracy and generalizability of our model. The ensemble model is similar to the algorithm we used to build the INT model, which is based on k-fold bagging. The final prediction model was the average of 50 ridge regression models (k = 5, 10 repetitions), and the choices for the regularization parameter were 21 values evenly distributed in a logarithmic scale, ranging from 0.01 to 100. Similar to nested cross-validation, the choice of the regularization parameter was determined based on out-of-bag cross-validation, and thus it’s only based on the training data. For each single model in the ensemble, we bootstrapped *n* – 1 participants with replacement from the *n* – 1 training participants and trained the ridge regression model based on the bootstrapped sample. To further increase the diversity of models in the ensemble, each time a participant was chosen by a bootstrapped sample, we also bootstrapped 4 runs with replacement from the participant’s data, and the face-selectivity map used in the regression was the average of the 4 bootstrapped runs. After all *n* – 1 participants had been chosen for the bootstrapped sample, we concatenated their vertices, and trained a ridge regression model based on the concatenated data. We obtained an estimated *S*_(*n*–1, faces)_ for each bootstrapped sample (coefficients of the regression model), and the final estimation of *S*_(*n*–1, faces)_ was the average across all bootstrapped samples. The model-predicted map of the left-out test participant was simply the matrix multiplication of the estimated stimulus descriptors *S*_(*n*–1, faces)_ based on the n – 1 training participants and the estimated tuning matrix *T_(p)_* of the test participant:

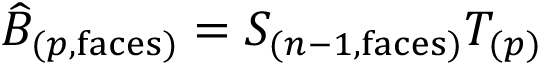

#### 4.6.3 Evaluating model-predicted maps

We evaluated the quality of model-predicted maps in the same way as (Jiahui et al., 2020). That is, for each test participant, we computed the Pearson correlation between the localizer-based map and the model-predicted map of the participant. Note that we estimated the reliability of the localizer-based map using Cronbach’s alpha, which is the expected correlation between two average maps, each based on 4 runs of independent data. Based on the Spearman–Brown prediction formula, we can estimate how Cronbach’s alpha changes with the amount of data (i.e., the number of localizer runs), and correspondingly, how much localizer data is needed to achieve the quality of the model-predicted map.

We also evaluated the specificity of our model-predicted maps. For each test participant, we also computed the correlations between the participant’s own localizer-based map and model-predicted maps of other participants. If the model-predicted map is highly specific to the participant, we expect the between-subject correlations to be much lower than the correlation with the participant’s own model-predicted map.

### 4.7. Predicting retinotopic maps

#### 4.7.1. Estimating retinotopic maps based on localizers

The *Forrest* dataset contains 4 retinotopic scans per participant that are 3 minutes each. The 4 runs are expanding rings, contracting rings, clockwise wedges and counterclockwise wedges, respectively. We followed the steps of (Warnking et al., 2002) and estimated an eccentricity map based on the runs of expanding rings and contracting rings and a polar angle map based on clockwise wedges and counterclockwise wedges for each participant. Specifically, we performed Fourier transformation on the time series data that were collected during stimulus presentation (5 cycles of 16 TRs [32 seconds] each; 80 TRs [160 seconds] in total; started 4 seconds after scan onset) and located the frequency component that had the same period as the stimuli (i.e., 5 cycles in 80 TRs). The amplitude of the component indicates to what extent a vertex’s response time series can be explained by retinotopic stimuli, and the phase of the component indicates the eccentricity or the polar angle that a vertex responds maximally to. Considering the hemodynamic response function of BOLD signal, we shifted the phase by 5 seconds to account for hemodynamic delay. For each kind of retinotopic map (i.e., eccentricity and polar angle), we averaged the Fourier transformation results of the two corresponding runs (e.g., expanding and contracting rings for eccentricity map) to get the final map. The amplitude was the mean amplitude of the two runs, and the phase was the circular mean of the two runs (which removes the remaining effects of hemodynamic delay).

#### 4.7.2. Model-predicted retinotopic maps

Each retinotopic map comprises two parts, namely an amplitude map and a phase map.

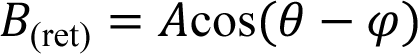

Here *A* is the amplitude, *θ* is the preferred phase (i.e., eccentricity or polar angle) for each vertex, and *φ* is the phase corresponding to the current stimulus. A vertex responds maximally when the phase of the current stimulus corresponds to its preferred phase, and the response decreases when the phase moves away from the vertex’s preferred phase. The retinotopic map can be modeled as a weighted sum of a sine map and a cosine map.

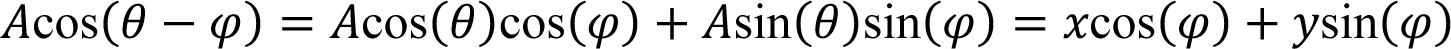

Note that the original phase *θ* is a circular variable and it’s difficult to predict it using a linear model (e.g., the model we used to predict category-selectivity maps). After the transformation, we have two new variables *x* and *y*, which contains the same information as the original amplitude map *A* and the phase map *θ*. However, both *x* and *y* are weights of the linear combination, and thus they can be predicted directly using linear models.

We used similar prediction procedures as the category-selectivity analysis for the current analysis. Specifically, we used leave-one-subject-out cross-validation, and the prediction models were ensembles of ridge regression models. For each test participant and each kind of retinotopic map, we trained two sets of ensemble models, one for predicting the weight map *x*, and the other for predicting *y*. After estimating the stimulus descriptors for *x* and *y* based on the training participants, we multiplied them by the estimated tuning matrix of the test participant to get the estimated *x* and *y* maps for the test participant. The model-predicted amplitude and phase maps can be computed from the estimated *x* and *y* maps:

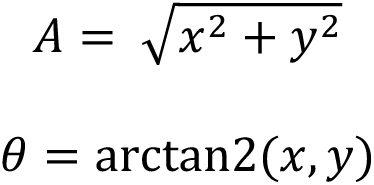

#### 4.7.3. Evaluating model-predicted maps

We evaluated the amplitude map and the phase map separately for each kind of retinotopic map. For the amplitude map, we computed the correlation between the test participant’s localizer-based map and the participant’s own model-predicted map, as well as the correlations with others’ model-predicted maps. We also computed Cronbach’s alpha based on the amplitude maps from the two runs from each kind of retinotopic map. In general, the amplitude maps were assessed in a similar way as the category-selectivity maps.

For the phase map, we computed the average (absolute) phase difference between the test participant’s localizer-based map and the participant’s own model-predicted map in the early visual cortex—an area known to have retinotopic responses. The early visual cortex was located based on regions V1, V2, V3, and V4 of the Glasser parcellation (Glasser et al., 2016). Similarly, we computed the average phase difference with others’ model-predicted maps, and the average phase difference between the two runs for each kind of retinotopic map. Note that the phase differences between the two runs are driven by both hemodynamic delay and noise, and their influences cannot be fully separated based on the current data.

### 4.8. Predicting response patterns to the movie

The previous analyses demonstrate the power of our model in predicting brain responses to new stimuli, such as object categories and retinotopic localizers. However, both object-category representation and retinotopy correspond to relatively coarse-grained cortical topographies. To assess the spatial granularity of our model, we further tested how well it could predict fine-grained spatial response patterns, such as time-point-by-time-point responses to a movie.

#### 4.8.1. Cross-validation scheme

For each movie dataset, we used leave-one-subject-out cross-validation to assess the model predictions. Each time, we built a template based on the full movie data of the *n* – 1 training participants. Similar to the distinctiveness analysis, we estimated the test participant’s tuning matrix using only half of the test participant’s movie data, and in this case it’s the first half of the movie data. The second half of the test participant’s movie data was held out for test. Then we multiplied the stimulus matrix for the second part of the movie with the estimated tuning matrix of the test participant to get the model-predicted response patterns to the second part of the movie that are based on other participants’ responses. We assessed the model prediction by comparing the measured response patterns and the model-predicted responses patterns of the test participant. Note that unlike our previous methods, in which we compared a participant’s response patterns to others’ patterns in the common model space, our INT model allows this comparison to be made in the native anatomical space (normalized to the fsaverage template) of each individual participant’s brain.

#### 4.8.2. Dimensionality reduction

For each time point (i.e., each TR), the response pattern is a vector of 18,742 elements. Similar to our previous work (Guntupalli et al., 2018, 2016; Haxby et al., 2011), we performed dimensionality reduction using principal component analysis (PCA) and compared the similarity of response patterns based on normalized PCs. We repeated the analysis using different numbers of PCs, ranging from 10 to 300 with an increment of 10. Note that the key results of this analysis (Figure 5D and 5E) are very robust against the choice of the number of PCs.

#### 4.8.3. Similarity between measured and predicted patterns

To illustrate the similarities of measured and predicted response patterns, we computed the correlations between measured and predicted response patterns based on 150 PCs. Specifically, we computed the similarities of patterns from the same participant and those from different participants; we also computed similarities of patterns for the same time point and those for different time points. These allowed us to evaluate the specificity of the model-predicted response patterns both to the participant and to the time point. Examples of the similarities are shown in Figure 5A and 5B, and the similarity distribution for each of the four conditions are summarized in Figure 5C.

#### 4.8.4. Binary movie time point classification

For each test participant, the similarity between the measured and predicted patterns for the same time point was much higher than those from different time points. We assessed to what extent this difference in similarity could be used to predict which time point of the movie the participant was viewing based on a binary classification task. The binary classification task is a 2-alternative forced choice. For each time point of the movie, we computed the correlation of its measured response pattern to two other response patterns—one was the pattern predicted from other participants’ responses to the same time point, and the other was the pattern predicted from other participants’ responses to another time point. The classification was successful if the similarity of patterns of the same time point was higher than the different time point, and thus the chance accuracy is 50%. We looped through all choices of the test time point, and for each test time point, looped through all choices of the foil time point and averaged the accuracies. Note that the difficulty of the binary classification task doesn’t change with the length of the movie data, and its accuracy can be considered as a measure of effect size in that sense. For example, the binary classification accuracy based on a dataset with 500 time points and another with 1000 time points are comparable. To evaluate the specificity of the predicted patterns to the test participant, we replaced the test participant’s predicted patterns with another participant’s predicted patterns and repeated the analysis.

#### 4.8.5. Multiclass movie time point classification

The classification accuracy of the binary classification task was close to 100%. To demonstrate the accuracy and specificity of the response patterns predicted by the INT model, we performed a multiclass movie time point classification analysis. That is, we compared the measured response pattern to a time point of the movie to all the model-predicted response patterns (i.e., predicted response patterns to all time points). We examined whether the pattern similarity was highest for the model-predicted response pattern of the same time point. The second part of the movie contains 1818 time points in total for the *Forrest* dataset, and 1680 time points for the *Raiders* dataset. Therefore, the number of choices was over 1000 for both datasets, and the chance accuracy was less than 0.1%. Note that the foils also included the time points right before or after the target time point, which was only 2 seconds (*Forrest*) or 1 second (*Raiders*) apart, and the inclusion of these neighboring time points made the classification task even more challenging.

### 4.9. Model performance with less data

In practice, it is not always feasible to collect a large amount of fMRI data during movie-watching as the datasets used in the current study (*Forrest*: 120 minutes; *Raiders*: 56 minutes). To assess the performance of our INT model with smaller data volume, we trained the model with smaller amounts of movie data for the test participant and evaluated its performance as a function of data volume.

First, we assessed how data volume affected the distinctiveness of the tuning matrix. This analysis requires two estimates of the same tuning matrix based on independent data, and thus each estimate can use up to half of the movie data (*Forrest*: 60 minutes; *Raiders*: 28 minutes). For the *Forrest* dataset, we repeated the analysis with 5, 10, 15, 20, 30, 40, 50, and 60 minutes of movie data for each estimate. For the *Raiders* dataset, we repeated the analysis with 5, 10, 15, 20, and 28 minutes of movie data for each estimate.

Second, we assessed how data volume affected the distinctiveness of local neural tuning based on a searchlight analysis. The same amounts of movie data as the whole-brain distinctiveness analysis were used. Instead of focusing on the average across searchlights, we assessed the 50^th^, 80^th^, 90^th^, 95^th^, and 99^th^ percentiles of the distribution.

Third, we assessed how data volume affected the estimation of category selectivity maps and retinotopic maps. Note that the objective of the analysis is to predict responses to new stimuli, and thus up to the entire movie data can be used to train the INT model and estimate the tuning matrices. For the *Forrest* dataset, we repeated the analysis with 5, 10, 15, 20, 30, 40, 50, 60, and 120 minutes of movie data. For the *Raiders* dataset, we repeated the analysis with 5, 10, 15, 20, 28, and 56 minutes of movie data.

Fourth, we used movie time point classifications to assess how data volume affected the quality of predicted response patterns to the movie. For this analysis, we used the same test data to evaluate the model, which was the second half of movie data for the test participant. Therefore, the movie data used to estimate the tuning matrix of the test participant was the first half of movie data or part of the first half. For the *Forrest* dataset, we repeated the analysis with 5, 10, 15, 20, 30, 40, 50, and 60 minutes of movie data. For the *Raiders* dataset, we repeated the analysis with 5, 10, 15, 20, and 28 minutes of movie data.

## Acknowledgments

We thank Caterina Gratton, Janine Bijsterbosch, and Pinglei Bao for helpful comments on earlier versions of the work.

This work was supported by NSF grants 1835200 (MIG) and 1607845 (JVH).

## Author Contributions

Conceptualization: MF, YOH, JVH; Methodology: MF, GJ, YOH, JVH; Software: MF, GJ; Formal analysis: MF; Data Curation: MF, SAN; Writing - Original Draft: MF; Writing - Review & Editing: MF, SAN, GJ, MIG, JVH; Visualization: MF; Resources: MIG, JVH; Supervision: MIG, JVH; Funding acquisition: MIG, JVH.

## Data and Code Availability

All data and code will be made available through DataLad (https://www.datalad.org/) and MF’s GitHub repository (https://github.com/feilong) upon publication. The *Forrest* dataset is also openly available through studyforrest (https://www.studyforrest.org/).

