## Supplementary figures for "The Individualized Neural Tuning Model: Precise and generalizable cartography of functional architecture in individual brains"

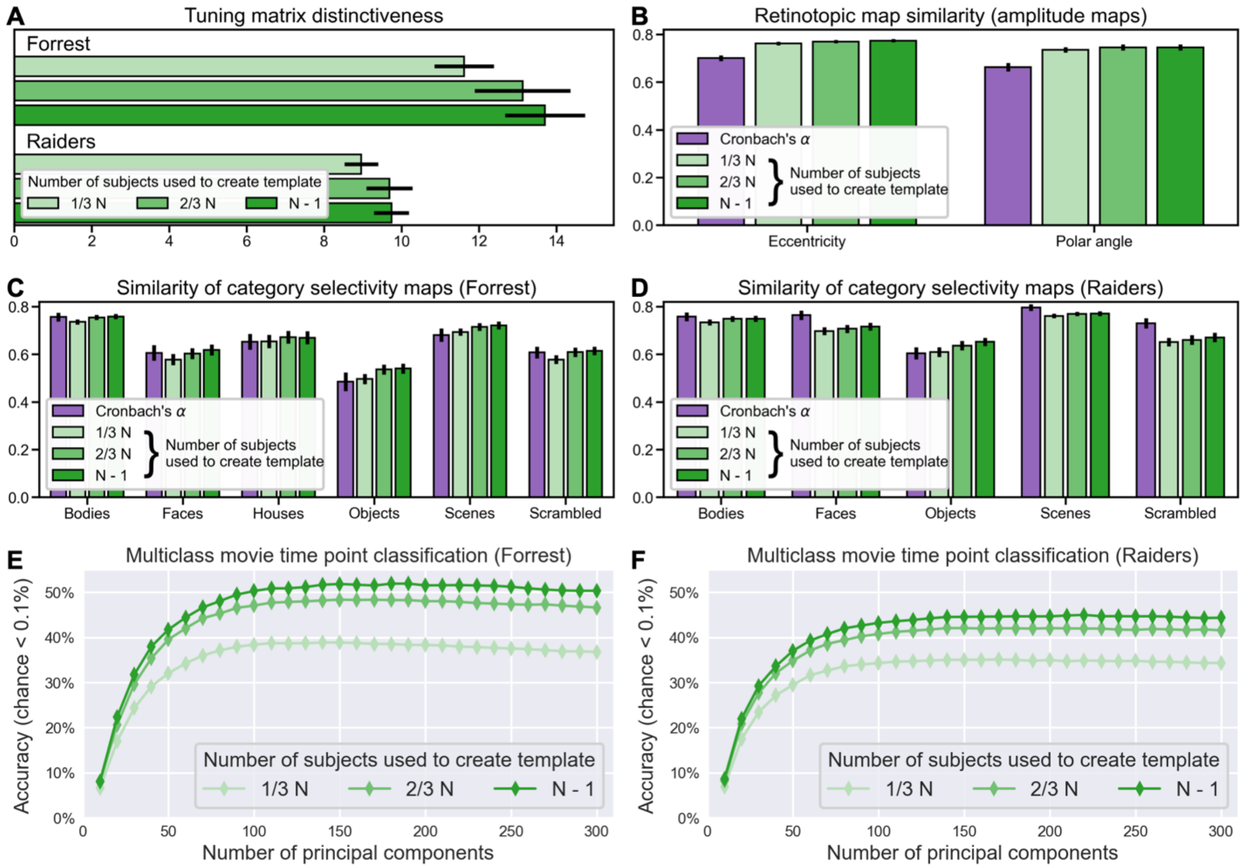


**Figure S1. Template quality as a function of the number of training participants.** The first step of our INT model re-represents each individual's neural response data matrix as a functional template linearly transformed with an idiosyncratic transformation, and thus the quality of the functional template may affect the performance of our INT model. We systematically manipulated the number of the participants used to create the functional template (i.e., training participants) and evaluated the performance of our INT model as a function of the number of training participants. We found that the performance of our INT model consistently increased with more training participants used to create the functional template. We observed the performance increase for the distinctiveness index **(A)**, the prediction of retinotopic maps **(B)** and category selectivity maps **(C–D)**, and the prediction of response patterns to movie time points **(E–F)**. This suggests that future development of the INT model may benefit from building the functional template based on more training participants than the current study (*Forrest*: *N* = 15; *Raiders*: *N* = 23). Error bars denote the standard error of the mean.

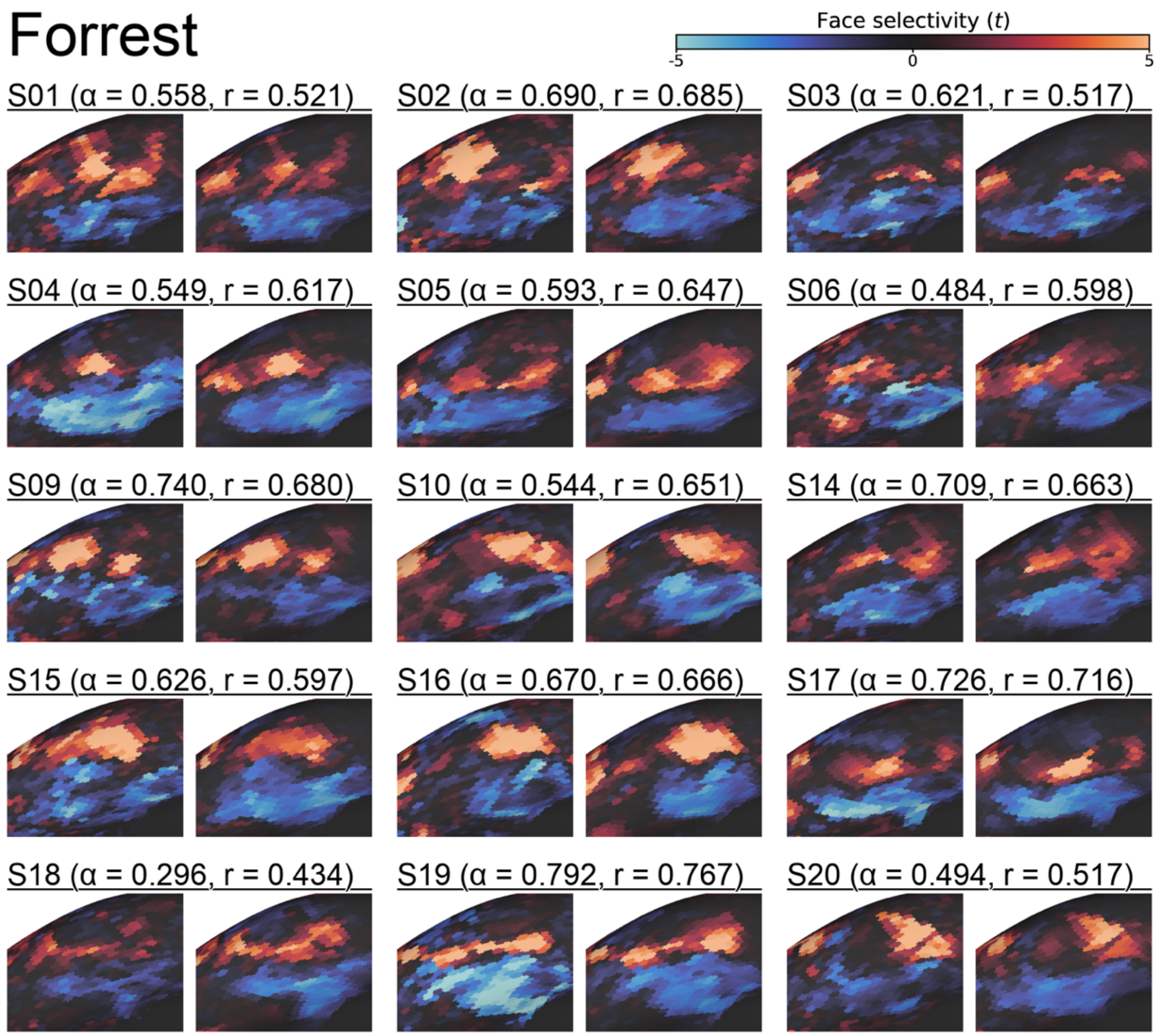


**Figure S2. Face selectivity maps for all *Forrest* participants.** For each participant, the estimated face selectivity map based on localizer scans is shown on the left, and the estimated map based on the INT model is shown on the right. Similar to Figure 3, the zoomed-in view of the right ventral temporal cortex is shown. α is Cronbach's alpha coefficient for the localizer-based maps, and r is the Pearson correlation between localizer-based and model-predicted maps. Both α and r are based on whole-brain face selectivity maps.

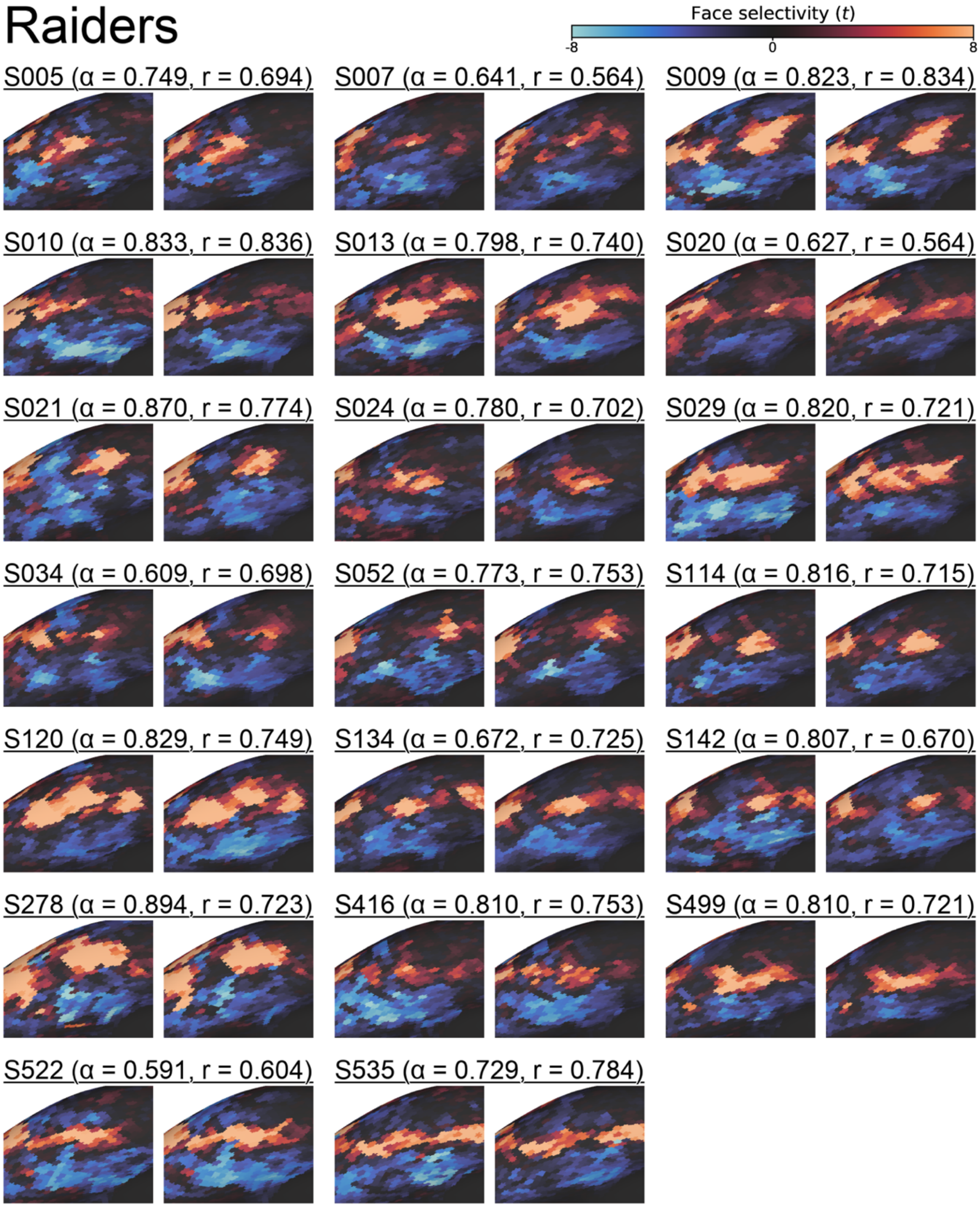


**Figure S3. Face selectivity maps for all *Raiders* participants.** For each participant, the estimated face selectivity map based on localizer scans is shown on the left, and the estimated map based on the INT model is shown on the right. Similar to Figure 3, the zoomed-in view of the right ventral temporal cortex is shown. α is Cronbach's alpha coefficient for the localizer-based maps, and r is the Pearson correlation between localizer-based and model-predicted maps. Both α and r are based on whole-brain face selectivity maps.

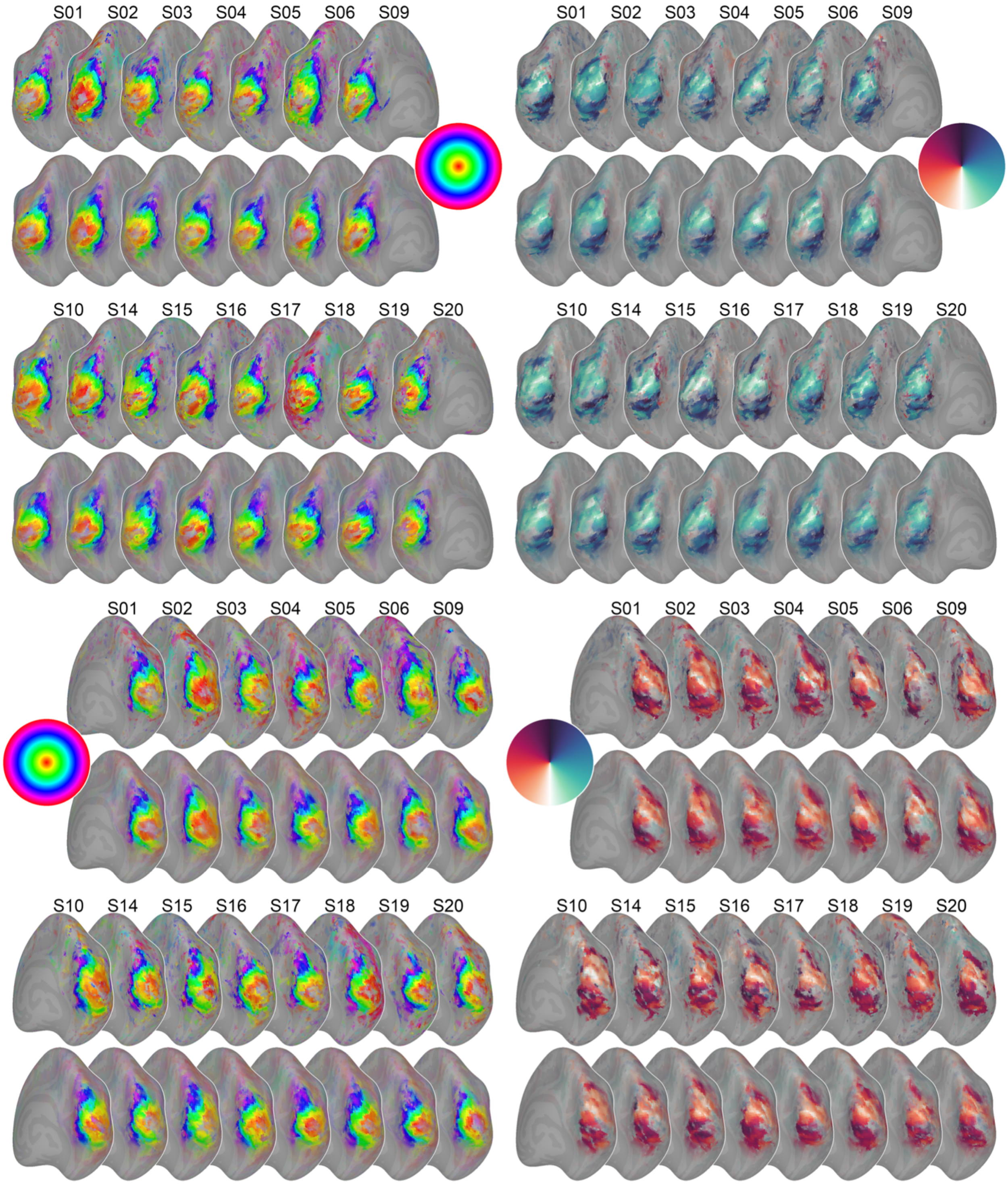


**Figure S4. Estimated retinotopic maps for all participants**. For each participant, the localizer-based map is shown in the top row, and the model-predicted map is shown in the bottom row. The retinotopic eccentricity maps are on the left side, and the polar angle maps are on the right side.
